# Geometric Quantification of Cell Phenotype Transition Manifolds with Information Geometry

**DOI:** 10.1101/2023.12.28.573500

**Authors:** Miao Huang, Yuxuan Wang, Junda Ying, He Xiao, Haijun Zhou, Lei Zhang, Weikang Wang

## Abstract

Cell phenotype transition (CPT) is crucial in development and other biological processes. Advances in single-cell sequencing reveal that CPT dynamics are confined to low-dimensional manifolds, yet current methods cannot directly quantify these manifolds. We present SCIM (single cell information manifolds), a geometry-guided approach using information geometry. SCIM embeds single-cell gene expression profiles as Gaussian distributions, defining the Fisher metric naturally in this space. We compute each cell’s coarse Ricci curvature, finding that low-curvature cells mark critical transitions. By further calculating each cell’s information velocity from RNA velocity, we find that regions with high information velocity coincide with low curvature, suggesting that geometry guides cellular dynamics on CPT manifolds. SCIM uncovers invariant features of CPT manifolds and provides a general framework for quantifying dynamics on these manifolds.

## Introduction

Biological systems, despite their inherent complexity, exhibit remarkable precision in various biological processes. A prime example is observed in embryo development, where the transition of stem cells into distinct phenotypes orchestrates the formation of well-organized organisms. Waddington’s landscape analogy is a well-known metaphor for illustrating embryo development. Recent advances in single-cell RNA sequencing have enabled a quantitative description of the landscape [1-3]. Most current methods define scalar potentials and fall short in fully capturing the dynamics of development, as sequencing is destructive to cells. To tackle this issue, algorithms have been developed to infer RNA velocity using counts of spliced and unspliced RNA, facilitating the learning of a vector field in the gene space of individual cells [4-8].

It is widely recognized that dynamics of cell phenotype transition (CPT) is confined to low-dimensional manifolds in phase space, which are governed by the underlying gene regulatory networks [5, 9-12]. However, current manifold learning methods merely preserve topology information in the original space and are often utilized as dimension reduction techniques that project high-dimensional data to a low-dimensional embedding space [13-16]. For instance, the coordinates for projecting scalar potentials or RNA velocity vectors with t-SNE or UMAP for visualization lack biological meaning and the capacity for quantitative study[17, 18]. Another major category of methods for learning the underlying manifold is probabilistic embedding. This category includes techniques such as diffusion maps, variational autoencoders (VAE), and various methods derived from these approaches [19-25]. Among these methods, scVI has been developed to address tasks such as batch correction, visualization, and clustering in single-cell RNA sequencing datasets [22]. Additionally, siVAE can identify gene modules and hubs by embedding cells and features simultaneously [23]. Furthermore, an integrated pipeline for deep probabilistic analysis of single-cell datasets, known as scvi-tools, is now available online [24].

In mathematics, Gauss and Riemann introduced the concept of intrinsic geometry of manifold, where the Riemannian metric determines the structure of manifold. Defining a Riemannian metric on the low-dimensional manifold with real data is challenging. Fisher information is a quantitative measure of the information that observables carry about parameters of probability distributions [26]. Rao demonstrated that Fisher information can be utilized as a Riemannian metric in probability space [27]. Amari developed information geometry based on the Fisher metric [27, 28]. From the perspective of the master equation, the dynamics of CPT can be viewed as the evolution of a probability distribution in phase space. By converting single cell gene vectors into probability distributions, the Riemannian metric can be established with Fisher information. One simple strategy for approximating the probability of single cells’ gene vectors is Gaussian mixture model (GMM), where each Gaussian distribution component represents the probability density of a stable cell state [29, 30]. For instance, Toshiaki Yachimura et al. developed a framework called scEGOT to infer cell differentiation dynamics using entropic optimal transport between GMM [31].

However, GMM is a coarse-graining approach that is incapable of yielding smooth manifolds. To overcome this limitation, we employ the Gaussian embedding method, in which each individual cell is modeled as a multivariate Gaussian distribution. This enables us to calculate the Fisher information of each cell and define the Riemannian metric in a smooth and continuous manner [32, 33]. The dynamics on the manifold are also important. Information velocity, which measures the rate of change in the probability distribution, can be used to quantify the speed of cell state variation on this manifold [34-36]. Information velocity can be computed based on the Fisher information metric and RNA velocity. Consequently, RNA velocity can be incorporated into the manifold formalism through the calculation of information velocity.

In this work, we introduce SCIM (single cell information manifolds), an approach to quantitatively analyze low-dimensional manifolds from single cell data. Initially, we transform each single cell’s sequencing data into a multivariate Gaussian distribution, calculate the Fisher information of each cell and quantify the manifolds of CPT [32, 33]. By representing each cell as a Gaussian distribution, we can capture its uncertainty and calculate the information distance between individual cells explicitly [26, 37]. Moreover, we can assess the Fisher metric on pre-defined coordinates such as eigengenes (EG), using the coordinate transformation with backward propagation in neural networks. By mapping the manifold onto coordinates with biological meaning, we can uncover the relationship between critical characteristics of cell dynamics and gene regulations. Unlike previous analyses, we concentrate on the invariant properties of the CPT manifolds, such as information velocity [34, 35, 38, 39]. Through incorporating information geometry with single cell geonomics analyses, SCIM provides a new framework for studying single cell transition dynamics, which can be further extended more rigorously from the perspective of geometry.

## Results

### Overview of SCIM

In this study, we harness neural networks to transform high-dimensional single cell data into a low-dimensional probability distribution space, represented with multivariate Gaussian distributions. To achieve this, we utilize a two-layer neural network in accordance with the Gaussian embedding method [32]. First, each single cell is represented as a vector *X*_*i*_ in the multidimensional expression space (Fig. 1a left), from which we compute the pairwise distance |*X*_*i*_ − *X*_*j*_| between cell *i* and cell *j*. Next, we use the k-nearest neighbor (kNN) cell graph (Fig. 1a right) to train a neural network (Fig. 1b top). The neural network *Φ*, which takes individual cell gene vector as input, comprises two hidden layers and simultaneous outputs mean (*µ*) and standard deviation (σ) of Gaussian distribution (*Φ*(*X*_*i*_) *= N*(*µ*_*i*_, σ_*i*_)). Cell triplets randomly sampled from the kNN cell graph (Fig. 1b bottom), serve as additional input alongside the cells’ gene vectors in training. We train the neural network with an energy-based learning method (Fig. 1b bottom) [40], where the energy is defined as the Fisher distance between the transformed Gaussian distributions and the loss function incorporates the graph structure information. The transformed Gaussian distribution captures the uncertainty of each cell node and reflects the diversity of its neighborhood (Fig. 1c) [32].

**Figure 1.**
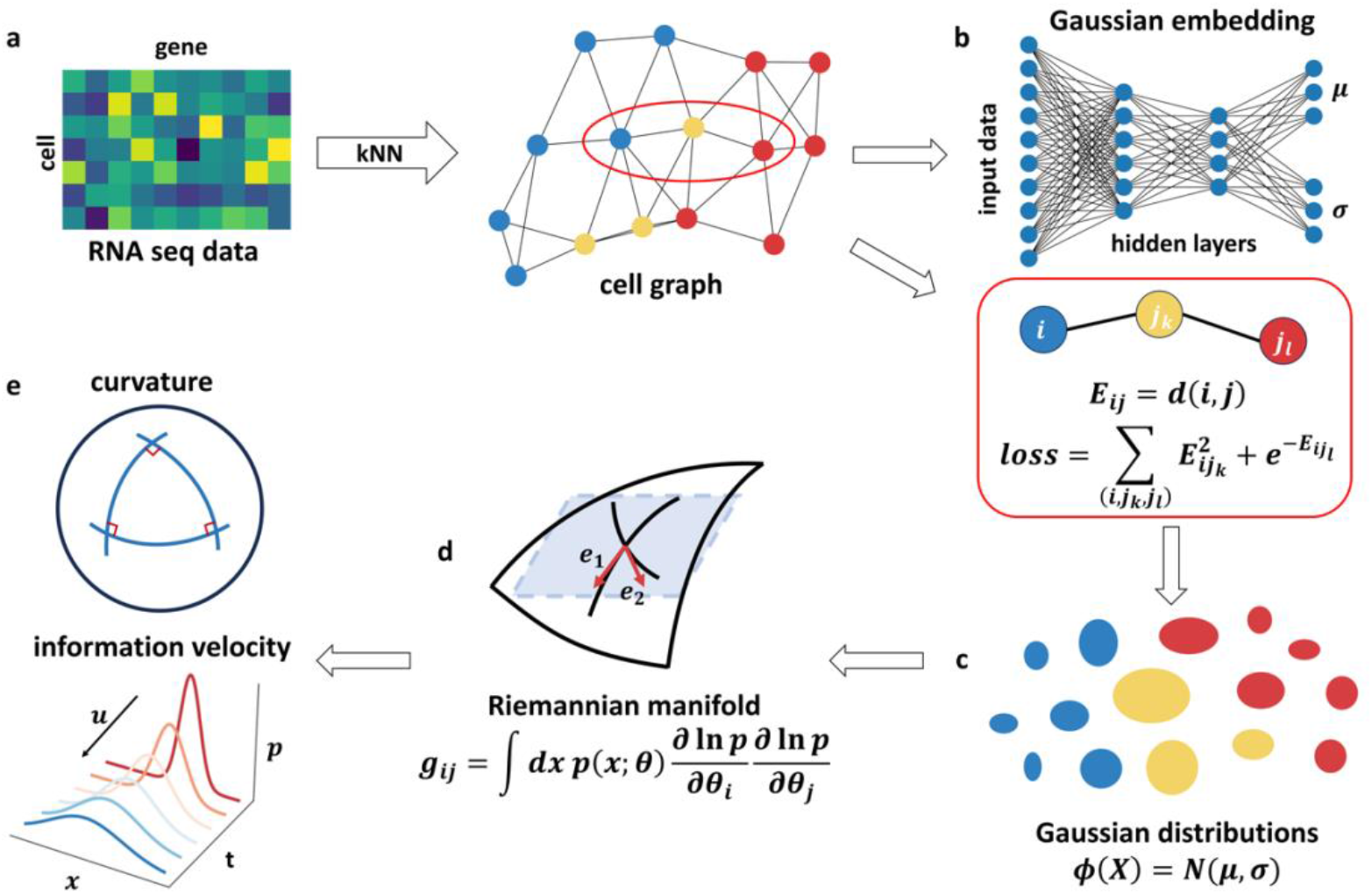
Illustrative overview of the SCIM framework. (a) Construction of kNN cell graph with single cell RNA sequencing data. Each single cell is depicted as a multi-dimensional vector, representing the counts of different genes’ RNA (left). The Euclidean distance between cells is calculated to build the graph. Colors represent different cell phenotypes. (b) The neural network architecture of the Gaussian embedding and its training method. The neural network is a multi-layer perceptron with two hidden layers. And it outputs *µ* and σ of Gaussian distribution respectively. Training is conducted using energy-based loss functions on sampled triplets. (c) Representation of single cells as multivariate Gaussian distributions. The transformed distributions are visualized as ellipses. And positions and sizes of the ellipses represent *µ* and σrespectively. (d) Defining Fisher metric on transformed Gaussian distributions give rise to information manifolds of CPT. In Gaussian distribution, parameters *θ* include both *µ*and σ. (e) The geometric property of CPT manifolds such as curvature and dynamics on the manifolds like information velocity are computed with SCIM.

By defining Fisher-Rao metric on these transformed Gaussian distributions whose parameters are *µ* and σ, the CPT manifolds can be quantitatively studied. The metric enables us to measure the distance on the manifold between different cells. Additionally, we can define other properties of the CPT manifolds like curvature and information velocity (Fig. 1d). Curvature, one of the most important geometric features on Riemannian manifolds, is approximated with a method developed by Yann Ollivier, bypassing the need to define a connection (Methods) [39].Information velocity, a property of dynamics, characterizes the variation rate of probability distribution and remains invariant under coordinate transformations (Methods).

In contrast to current manifold learning methods, which are tailored for visualization, our method, SCIM, is not constrained to this purpose. Although we can simply project high-dimensional cell vectors onto a 2D or 3D (*µ*, σ) space for visualization, it is crucial to recognize that such projections can misrepresent or distort the original manifold’s geometry. This is due to the inherent limitations of lower-dimensional representations in capturing the true structure of high-dimensional manifolds. Different from previous studies that rely on visualization, SCIM enables us to study the intrinsic geometry of the manifold quantitatively and identify invariant properties that are independent of coordinate settings.

### Application of SCIM on toy datasets

To evaluate the effectiveness of SCIM, we apply it to several test datasets. The first dataset consists of data points sampled on a saddle surface (Fig. 2a top). The second dataset consists of data points sampled on the surface of a 3D ellipsoid (Fig. 2a middle). We also generate a third dataset by sampling on a four-well potential surface through simulating multiple trajectories (Fig. 2a bottom). For each dataset, we add seven-dimensional random features (Gaussian distribution with small amplitude) as noise to form a ten-dimensional data, and construct kNN graphs from the generated samples. Using SCIM, we project the embedded data of saddle surface, ellipsoid and four-well potential surface on the plane of(*µ*_1_, *µ*_2_), (*μ*_1_, σ_1_)and (σ_1_, σ_2_) respectively (Fig. 2b). For comparison, we also apply current manifold learning methods, such as SNE, UMAP and diffusion map, to the test datasets (Fig. S1a).

**Figure 2.**
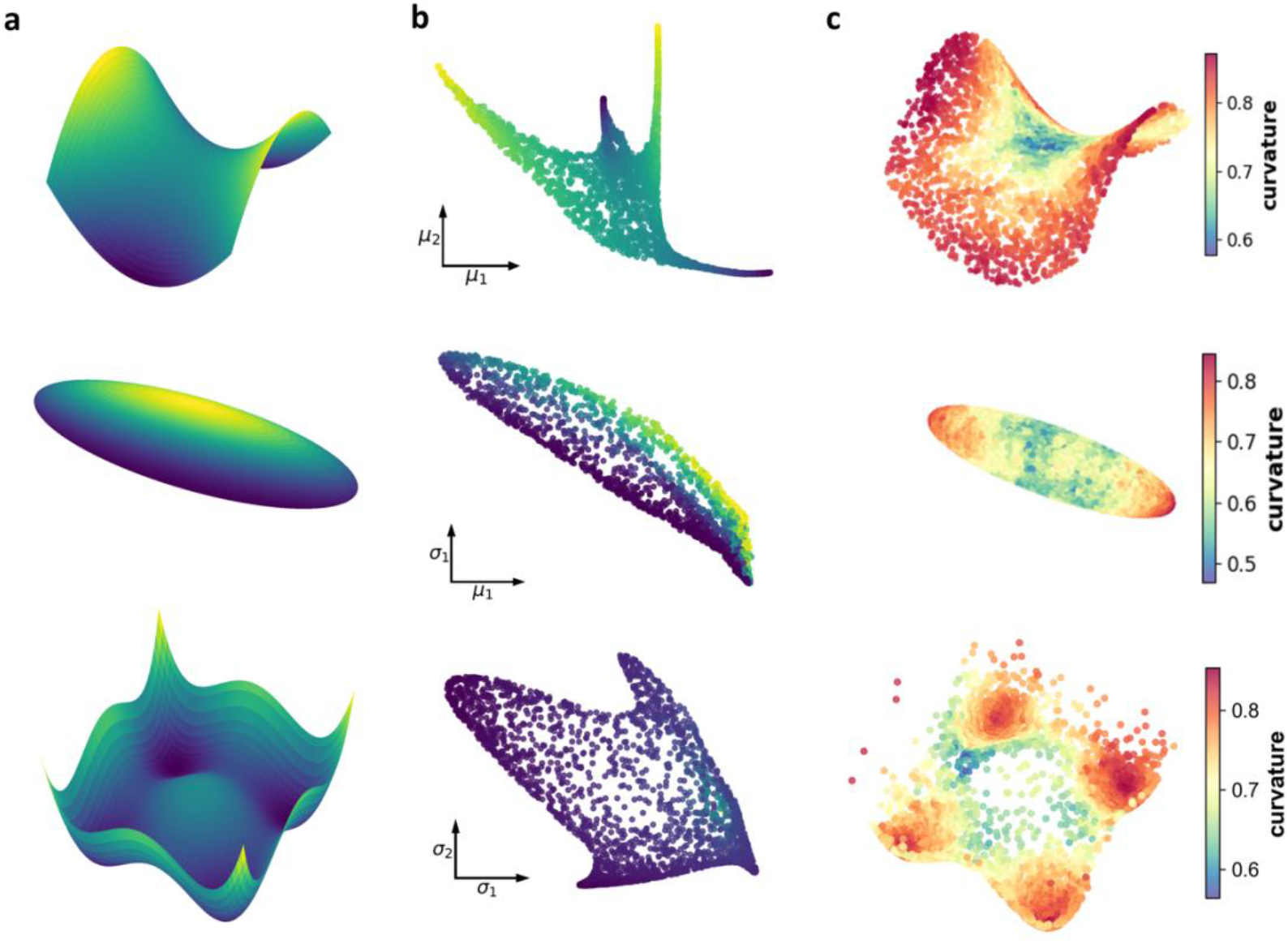
Evaluation of SCIM on toy models including saddle surface, ellipsoid and four-well potential. (a) Sampling of data points on various surfaces: saddle surface (top), ellipsoid surface (middle), and simulated trajectories on a four-well potential landscape (bottom). (b) Planar projections of saddle surface on the plane of (*µ*_1_, *µ*_2_)(top), ellipsoid on the plane of (*µ*_1_, σ_1_) (middle) and four-well potential on the plane of (σ_1_, σ_2_) (bottom) respectively. (c) CRC distributions across the sampled data points in saddle surface (top), ellipsoid surface (middle) and four-well potential (bottom). Color represents the value of CRC.

To evaluate the preservation of local metrics, we calculate ratios of geodesic lengths of multiple semi-circles sampled on a sphere surface before and after embedding, and compare the resulting ratio distributions between SCIM and other manifold learning methods (Methods). As manifold learning methods like t-SNE and UMAP primarily tend to preserve global topology for visualization purposes, their length distributions show larger coefficient of variation (CV) value comparing with that of SCIM. And diffusion maps and VAE embeddings also exhibit larger CV value in length distributions (Fig. S1b).

Curvature is a fundamental property in Riemannian geometry, but its calculation requires a defined connection, which can be challenging due to the limitation of data and requirement on resolution of difference methods. To circumvent this, we use coarse Ricci curvature (CRC) for estimating curvature on complex networks (Methods) [38, 39]. This is an approximation method that utilizing the relation of transportation distances between probability distributions and Ricci curvature. For the saddle surface data, data points near the saddle point have lower CRC values compared to other regions (Fig. 2c top). In the ellipsoid data, we observe that the data points at both ends of the major axis show higher CRC values (Fig. 2c middle). In the four-well potential landscape, data points on the barriers between basins have lower CRC values, consistent with the findings in the saddle surface data (Fig. 2c bottom). Previous studies on complex networks has shown that CRC values of edges between nodes in different communities are generally negative, whereas edges inside each community usually have positive CRC values [41]. Our results, in conjunction with prior research, reveal that saddle points or the intermediate region between communities display lower CRC value. We further validate the robustness of our method by calculating CRC on saddle surface with varying parameter settings (Fig. S2).

Additionally, in the four-well potential data, points in the basins show high CRC values (Fig. 2c bottom). An intuitive explanation is that data points within a basin share similar neighbors, leading to close embedded distributions, short Wasserstein distances and high CRC values. These insights from toy datasets pave the way for identifying meta-stable cell states and critical points in CPTs of single-cell RNA sequencing data.

We applied the VAE to all the toy datasets (Fig. S3a). Except for the ellipsoid data (Fig. S3b middle), the distributions of CRC differ from those obtained with SCIM (Fig. S3b). A major problem with VAE is that the CRC results are not self-consistent. In the saddle surface data, not only do the points around the saddle points exhibit low curvature, but the entire ridge does as well (Fig. S3b top). Conversely, in the four-well potential, the barriers exhibit high curvature (Fig. S3b bottom). Additionally, the CRC value is significantly affected by the relative weight of the KL-divergence term in the loss function (Fig. S3c).

### Fisher information analyses of single cell RNA sequencing data

We first applied SCIM to single-cell RNA sequencing data of dentate gyrus neurogenesis (Fig. 3a). Throughout the process of neurogenesis in dentate gyrus, radial glia-like cells differentiate through neuronal intermediate progenitor cells (nIPCs), neuroblast 1 and 2, immature granule cells, and eventually mature into granule cells [42].

**Figure 3.**
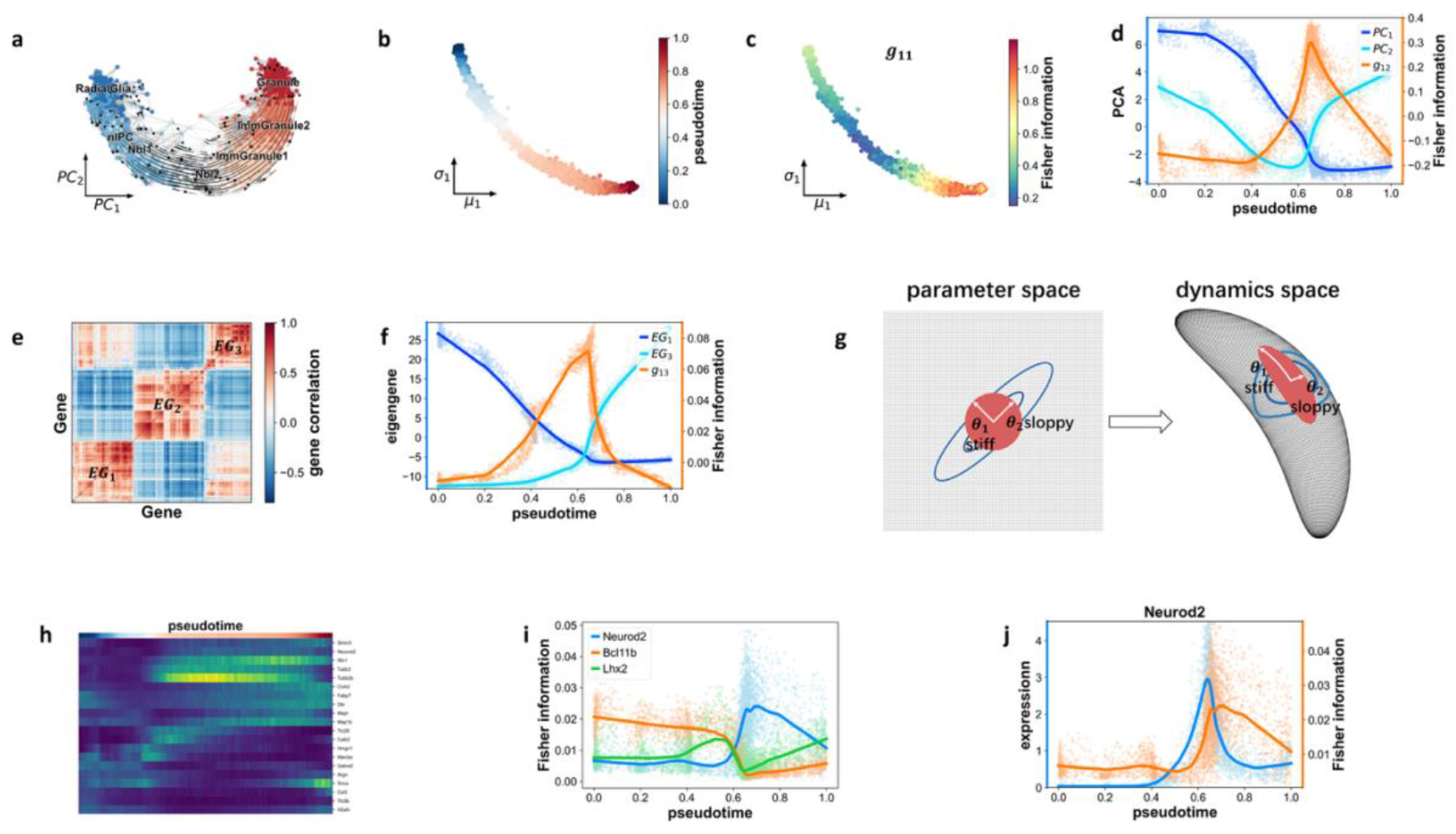
Fisher information analyses of dentate gyrus neurogenesis. (a) KNN graph of dentate gyrus neurogenesis with RNA velocity flow projection. Color represents cell type. The direction of transition denoted by the flow is from radial glia cells to granule cells. (b) Pseudo-time analysis of dentate gyrus neurogenesis (projection on the plane of(*µ*_1_, σ_1_). Color represents pseudo-time. (c) Variation of *g*_11_ (Fisher information of*µ*_1_) in dentate gyrus neurogenesis. Color represents value of Fisher information. (d) Variation of one non-diagonal element in Fisher information matrix (*g*_12_) on PC coordinates (PC1-PC2) and values of PC1 and PC2 (with respect to pseudo-time). This non-diagonal element exhibits a peak during the transition process. The lines are fitted with LOWESS (Methods). (e) Eigengenes (EG) derived with clustering on gene correlations. Color represents correlations among different genes. The value of eigengene is defined as PC1 of each cluster. (f) Variation of one non-diagonal element in Fisher information matrix (*g*_12_) on eigengene coordinates (EG1-EG2) and the corresponding values of eigengenes (with respect to pseudo-time). This non-diagonal element also exhibits a peak during the transition process and position of the peak is close to that in PC coordinates in panel (e). The lines are fitted with LOWESS (Methods). (g) Schematic illustration of stiff and sloppy parameters. The system’s dynamics are determined by the parameters, shown as a mapping from the parameter space to the dynamics space. Variations in stiff parameters significantly impact the system’s dynamics, while the influence of sloppy parameters is negligible (red discs). Given some samples from the dynamics space, stiff parameters can be inferred with small uncertainties, whereas sloppy parameters are poorly determined (blue circles). (h) Heatmap of Fisher information for multiple genes with respect to pseudo-time. The variation in Fisher information reflects changes in the susceptibilities of different genes during the CPT process. (i) Fisher information of several transcriptional factors, including Neurod2, Bcl11b, and Lhx2, in each single cell with respect to pseudo-time. The variation in Fisher information reflects the different regulatory functions of these transcriptional factors during the CPT process. Each dot represents a single cell, and the lines are fitted using LOWESS (Methods). (j) Fisher information (orange) and expression value (blue) of Neurod2 in each single cell with respect to pseudo-time. Each dot represents a single cell, and the lines are fitted using LOWESS (Methods).

For visualization, we calculated the pseudo-time of each single cell and plot the single-cell data in the probability distribution space (Fig. 3b and Fig. S4a). We also showed the Fisher information (*g*_11_) of *µ*_1_ in Fig. 3c (Other elements in Fisher information matrix are shown in supplement Fig. S4b).

The coordinates (*µ*, σ) lack direct biological interpretation. To elucidate the relationship between the geometry of CPT manifold and the biological characteristics of the system, it is necessary to calculate the Fisher metric in different coordinate systems, particularly those with explicit biological meaning. Here we performed a coordinate transformation to obtain the Fisher metric:

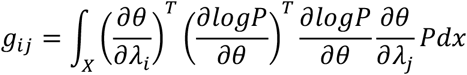

in the new coordinates λ. Here *θ* represent coordinates (*µ*, σ) (partial derivative 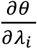 calculations are detailed in Methods). *i* and *j* are indexes of the new coordinates.

We first converted the (*µ*, σ) coordinates to principal component (PC) coordinates. The Fisher information is a powerful tool for analyzing informative orthogonal parameters [43]. With SCIM, we can directly obtain the diagonal Fisher matrix. However, the Fisher matrices in PC coordinates may not necessarily be diagonal after coordinate transformation. If a non-diagonal element *g*_*ij*_ (*i* ≠ *j*) of Fisher matrix equals zero, parameters λ_*i*_ and λ_*j*_ are defined as information orthogonal [43]. By analyzing the Fisher matrix in coordinates with biological meaning, we can discern the orthogonality between these parameters and draw some biological conclusions. As shown in Fig. 3d, the Fisher information of PC1 and PC2 is relatively low (around 0) most of time except for a peak at pseudo-time around 0.6, coinciding with drastic changes in PC1 and PC2 values.

We also performed coordinate transformation from (*µ*, σ) to eigengene coordinates, where eigengene represent the PC1 of gene modules identified through gene correlation clustering (Fig. 3e) (Methods). During neurogenesis, the non-diagonal element *g*_13_ in the Fisher matrix of eigengenes remains nearly zero all the time. This information orthogonality between both eigengenes reflects their distinct roles at different neurogenesis stages. Similar with that of PC coordinates, this non-diagonal element *g*_13_ also peaks at pseudo-time around 0.6 (Fig. 3f). This consistency between two different coordinate systems reflects the underlying regulatory mechanism independent of coordinates. The Fisher information and information orthogonality are dynamical as the Fisher matrix evolves over time. Through Fisher information analysis, we gain diverse perspectives on the dynamics of CPT process. It is important to note that Fisher information is affected by the choice of coordinates, for instance, the Fisher information of two gene modules’ PC2 behave differently from that of eigengenes (Fig. S4c).

Fisher information reflects the sensitivity of probability density to control parameters. Stiff parameters are those with large values of Fisher information, while sloppy parameters have small values (Fig. 3g) [44]. By calculating the Fisher information for each gene (Methods), we can identify the stiff and sloppy genes in the context of CPT. Our analyses reveal that the stiffness and sloppiness of genes vary throughout the CPT process, suggesting that different genes may play different roles at different stages of CPT (Fig. 3h). Specifically, the Fisher information of transcriptional factors reveals their probable regulatory functions in CPT. For instance, the Fisher information of Neurod2 against pseudo-time exhibits a peak around a pseudo-time of 0.6 (Fig. 3i & 3j), which aligns with the trends of non-diagonal Fisher information in PC coordinates (Fig. 3d) and eigengenes (Fig. 3f). In contrast, the Fisher information of Bcl11b and Lhx2 behaves differently (Fig. 3i). Therefore, Fisher information analysis provides a potential method for identifying the roles of different genes in regulating CPT.

We compared the embeddings of SCIM and scVI on dentate gyrus neurogenesis (Fig. S5). As shown in Fig. S5b, the scVI embeddings form spherical clouds around the origin in the coordinate system due to the regularization in VAE, which minimizes its Kullback-Leibler divergence (KL-divergence) with the standard normal distribution *N*(0,1) [20]. And the latent embedding in VAE is designed to facilitate generative sampling for the decoder. In contrast, SCIM focuses on preserving local metrics and provides a framework for quantifying geometric properties of the underlying low-dimensional manifolds.

We also apply Fisher information analysis on two additional datasets. One is the development of pancreatic endocrine cells (Fig. 4a) [45]. During embryonic development Ngn3-low progenitors first transform into Ngn3-high precursors, then Fev-high (Fev+) cells. The latter further develop into endocrine cells including glucagon producing α-cells and β-cells. Here we concentrate on the α-cell branch. The variation of Fisher information of *µ*_1_ is shown in Fig. 4b. And the Fisher information of the two eigengenes (non-diagonal element) exhibits one local minimum value and one local maximum value as CPT proceeds (Fig. 4c). And both local extrema are deviated from zero, indicating the complex interplay between eigengenes and their synergetic influences on the probability distributions vary as CPT proceeds. The variation in Fisher information for multiple genes during pancreatic endocrinogenesis reveals the switching of roles as stiff or sloppy genes throughout this CPT process (Fig. 4d). Additionally, the changes in Fisher information for transcriptional factors may indicate the dynamics of gene regulation at different stages (Fig. 4e). For example, the Fisher information and expression levels of Neurog3 are plotted against pseudo-time (Fig. 4f).

**Figure 4.**
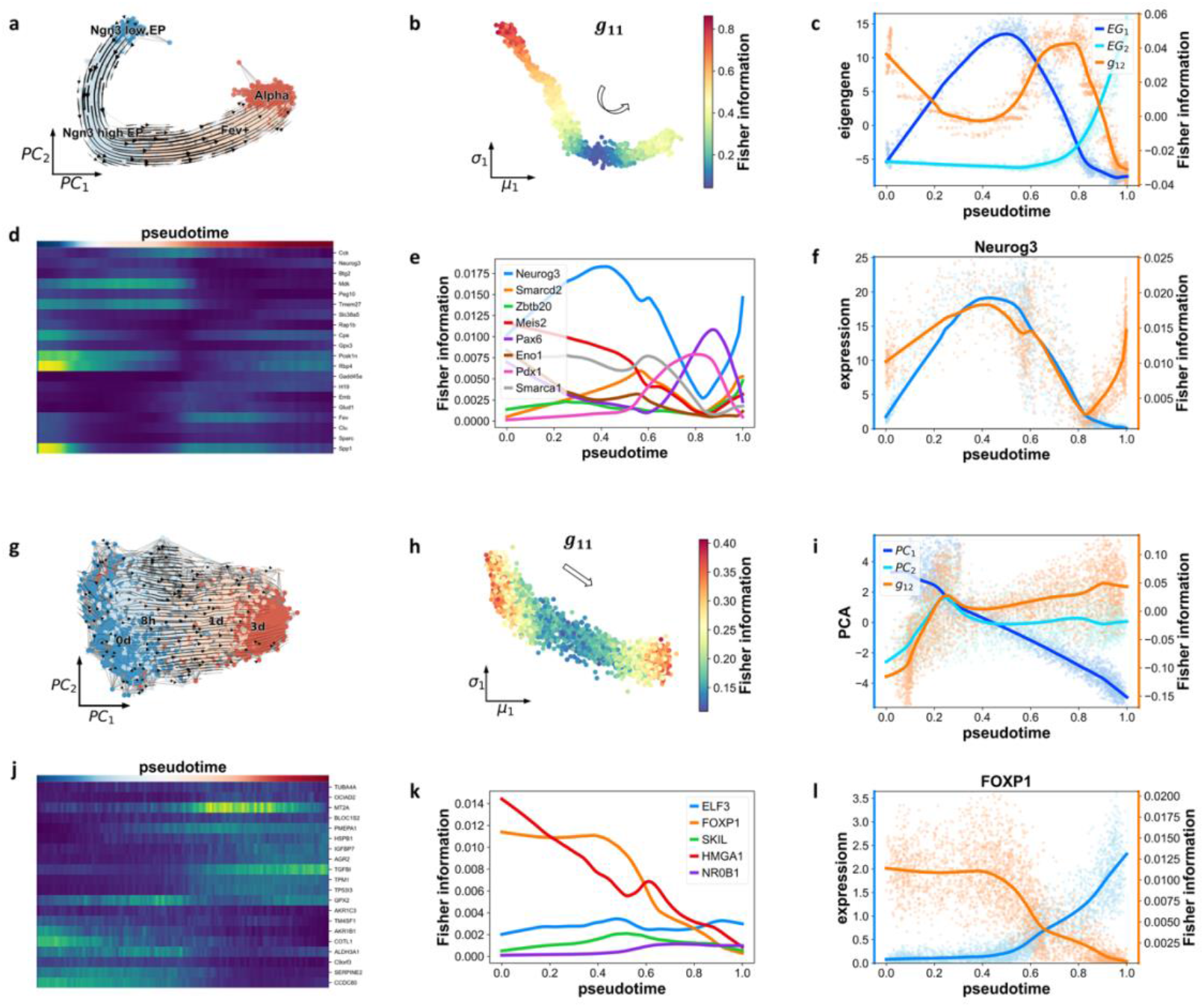
Fisher information analyses of pancreatic endocrinogenesis and EMT of A549 cells induced by TGF-β. (a) kNN graph of pancreatic endocrinogenesis with RNA velocity flow projection. Color represents cell type. (b) Variation Fisher information of (*µ*_1_) in pancreatic endocrinogenesis. Color represents value of Fisher information. (c) Variation of one non-diagonal element in Fisher information matrix (*g*_12_) on eigengene coordinates and the corresponding values of eigengenes (with respect to pseudo-time) in pancreatic endocrinogenesis. The lines are fitted with LOWESS (Methods). (d) Heatmap of Fisher information for multiple genes with respect to pseudo-time. The variation in Fisher information reflects changes in the susceptibilities of different genes during pancreatic endocrinogenesis. (e) Fisher information of various transcriptional factors in each single cell with respect to pseudo-time. The lines are fitted to single-cell data using LOWESS (Methods). (f) Fisher information (orange) and expression value (blue) of Neurog3 in each single cell with respect to pseudo-time. Each dot represents a single cell, and the lines are fitted using LOWESS (Methods). (g) Same as in panel (a), except for EMT of A549 cells. (h) Same as in panel (b), except for EMT of A549 cells. (i) Same as in panel (c), except for Fisher information of PC1-PC2 in EMT of A549 cells. (j) Same as in panel (d), except for EMT of A549 cells. (k) Same as in panel (e), except for EMT of A549 cells. (l) Same as in panel (f), except for FOXP1 in EMT of A549 cells.

The other is an induced in-vitro transition process, the epithelial-to-mesenchymal transition (EMT) of human A549 cells treated with TGF-β for different durations (0 day, 8 hours, 1 day, and 3 days) (Fig. 4g) [46]. At the initial stage of EMT (0 day), most A549 cells are in the epithelial state. And most cells transit into the mesenchymal state after three days of treatment. Compared with the development datasets, cancer cells exhibit greater heterogeneity, reflected in the significant Fisher information variation of *µ*_1_ in the 8h sample (Fig. 4h). The non-diagonal element of Fisher information matrix between two PCs is non-zero initially (mostly 0 day). And the two PCs become information-orthogonal after TGF-β treatment (Fig. 4i). Due to cell state heterogeneity, we calculate pseudo-time for a clearer dynamic representation. The changes in Fisher information for multiple genes (Fig. 4j) and transcriptional factors (Fig. 4k) reveal variations in control during the EMT process. We also presented the Fisher information and expression levels of FOXP1 across pseudo-time (Fig. 4l). It appears that there is no clear correlation between the magnitude of Fisher information and gene expression (Fig. 3j, 4f, and 4l).

The Gaussian embedding of each single cell can also be utilized for cell type classification. To achieve this, we developed a clustering method by integrating Gaussian embedding with agglomerative hierarchical clustering (Methods). When compared to other methods such as Leiden, Louvain, DTNE, and multiscale PHATE (ms-PHATE) algorithms (see Fig. S6) [47-50], our approach demonstrates similar accuracy in terms of the Rand Index (RI), Mutual Information (MI), and F1-score. This indicates that SCIM is capable of extracting key features related to cell states, thereby reinforcing the foundation for further analyses based on Fisher information.

### Identification of critical points with curvature

Analyzing Fisher information in specific coordinates can shed light on the regulatory dynamics in CPT. However, the variation of Fisher matrix is dependent on the coordinate system. To learn more about the geometry of the information manifold, we further computed CRC on the single-cell RNA sequencing datasets (Fig. 5a) (Methods).

**Figure 5.**
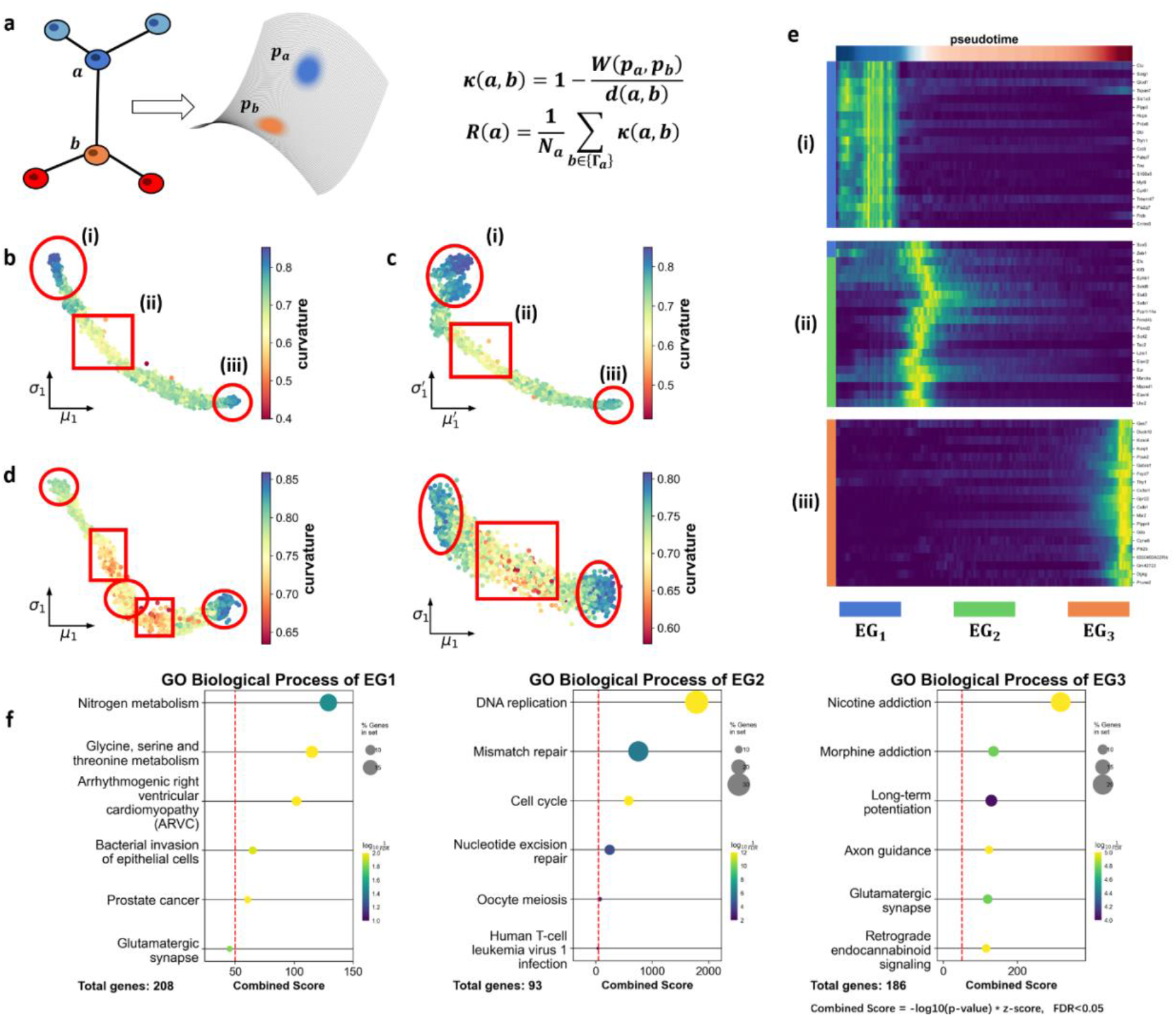
Identification of critical points and basins in different CPT processes with CRC. (a) Schematic representation of CRC calculation from single cell RNA sequencing data. Single cell gene vectors are transformed into Gaussian distributions. The ratio between Euclidean distance of cells and Wasserstein distance of transformed distributions is calculated to derive CRC. In the equation, *W*(*P*_*a*_, *P*_*b*_) represents the Wasserstein distance between the Gaussian embeddings of cell a and cell b and *d*(*a, b*) denotes the Euclidean distance between cell a and cell b. *k*(*a, b*) is the CRC of the edge between cell *a* and cell *b*, and *R*(*a*) is the CRC of cell *a*. {Γ_a_} represents all the neighboring cells of cell *a* and N_a_ is the number of neighbors of cell *a*. (b) Identification of the critical points and basins with CRC in dentate gyrus neurogenesis. Regions where cells have local extreme (maximum and minimum) values of CRC are enclosed in red box. High CRC values are found in regions (box i and iii) corresponding to radial glia cells and granule cells, whereas the intermediate region exhibits low CRC values (box ii) that corresponds to saddle node. The color bar is reversed in this panel for better visualization. (c) Stability of CRC in different Gaussian embedding coordinates. The distribution of CRC calculated within a different Gaussian embedding coordinate (*µ*^′^, σ′) is similar to that within (*µ*, σ) in panel (b). The color bar is reversed in this panel for better visualization. (d) Identification of the critical point and basins with CRC in pancreatic endocrinogenesis (left) and EMT of A549 cells induced by TGF-β (right). In the pancreatic endocrinogenesis, besides the high-CRC regions at the initial and final stages in transition, an additional high-CRC region is detected in the middle. Two regions with low CRC values (rectangles) bridge the three high-CRC regions (circles). In the EMT dataset, the low CRC cells are mainly found in the intermediate region (rectangles) between the 8-hour and 1-day samples. The color bar is reversed in this panel for better visualization. (e) Heatmap of genes correlated with CRC values. Genes correlated with CRC values are highlighted within the red boxes labeled i, ii, and iii. Adjacent color bars on the left side of the heatmap denote the eigengenes associated with each gene in the row. (f) Bubble chart of Gene Ontology (GO) enrichment analysis of different eigengenes in dentate gyrus neurogenesis. The red dashed line indicates the combined score threshold of 50. The False Discovery Rate (FDR) was calculated using the Benjamini-Hochberg method. The top six terms with FDR < 0.05 are shown for each eigengene. The size and color of the circle represent the proportion of significant genes in the total genes and the corresponding FDR value, respectively.

In the dentate gyrus data, cells with the lowest and highest pseudo-time values correspond to radial glia cells and granule cells respectively and these cells displayed high CRC values. As demonstrated in Fig. 2d, high-CRC value characterizes the basins on the energy landscape, indicating the two types of cells are in stable states. Following our finds on toy datasets, cells in this low-CRC region are posited to be around the saddle point, a critical point in dynamical system theory that connects different attractors or basins. And we do observe a low-CRC region (Fig. 5b box ii) bridges the two high-CRC regions (Fig. 5b box i and iii), suggesting a transition from radial glia cells to granule cells. Therefore, this low-CRC region is associated with the intermediate or transition states which often display mixed properties of different cell phenotypes and are typically unstable and revisable. In addition, the distribution of Ricci curvature should be consistent across different Gaussian embeddings. Here we use SCIM to transform the single cell data of dentate gyrus to another Gaussian embedding coordinates (*µ*′, σ′) (Fig.5c). We find that the CRC distribution is similar with that of (*µ*, σ) in Fig. 5b and one can still observe a low-CRC region (Fig. 5c box ii) bridges the two high-CRC regions (Fig. 5c box i and iii).

In the pancreatic endocriogenesis data, two low-CRC regions are identified, bridging three high-CRC regions (Fig. 5d left). And there is a subset of Ngn3-high precursors in the meta-stable state (high CRC region in the middle) between the two transitory regions with low-CRC values, suggesting a step-like transition process in endocriogenesis. For the EMT dataset, curvature analysis reveals that cells treated with TGF-β for one day have the lowest CRC value, connecting two high-CRC regions that are the epithelial and mesenchymal states (Fig. 5d right). The intermediate state is commonly recognized as a partial EMT state which is related to the collective migration of tumors [51, 52].

SCIM effectively identifies transition states characterized by low CRC values, revealing a connection between the geometric properties of CPT manifolds and their biological functions. We have observed that low CRC values arise from relatively large Wasserstein distances between neighboring cells, indicating a heterogeneous cell population. Conversely, in regions with high CRC values, cells are more homogeneous, as evidenced by shorter Wasserstein distances between neighbors (Fig. S7). This observed heterogeneity aligns with existing knowledge about transition states, where cells are exploring the state space [53].

Characterizing transition states in CPT is crucial for understanding cell fate decisions and interpreting early warning signals for disease transitions [53-57]. Various methods, including scGeom, scTite, and CellTran [58-60], have been developed to identify transition states. Notably, scGeom analyzes transition cells using CRC, similar to SCIM, and both methods consistently find that transition cells exhibit low CRC values [58]. Meanwhile, scTite utilizes transition entropy to identify transition cells [59]. And CellTran assigns a transition index based on the gene expression correlation between cells and their neighbors [60]. We compared SCIM with these methods using the dataset from this study (Fig. S8) and additional test datasets from the respective publications (Fig. S9). SCIM demonstrated consistent performance across all datasets. For instance, validation of SCIM on simulated data in CellTran and real datasets in scGeom demonstrated that SCIM’s CRC reliably corroborates findings from both CellTran and scGeom. While other methods either fail to identify transition cells in simulated data or mistakenly classify cells at terminal stages as transition cells, which contradicts the understanding that terminal states are generally stable rather than transitional.

For the dentate gyrus neurogenesis data, we analyzed genes that are highly correlated with CRC values in the marked regions (boxes in Fig. 5b&5c), i.e. the basins and critical point. These genes serve as markers of the corresponding cell regions. And they correspond to three eigengenes respectively (except a few genes in region ii belong to the first eigengene), highlighting the modular regulation in CPT (Fig. 5e). We further examined the biological functions of the eigengenes by performing gene ontology (GO) analysis on dentate gyrus neurogenesis data (Fig. 5f) (Methods) [61]. The GO terms highly associated with the eigengenes are listed. For example, the third eigengene (EG3) is closely associated with neural functions.

It is important to note that CRC is an approximation of Ricci curvature and is not derived from Fisher metric. Nevertheless, our analyses reveal its association with the metric. The low-CRC region coincides with the peaks of non-diagonal element of Fisher information matrix (Fig. 3d & 3f).

### Geometry guides single cell dynamics on CPT manifolds

CRC elucidates the structure of the CPT manifold, with low-CRC regions closely related to saddle points between basins and its derivation does not require dynamics information. In the spirit of general relativity theory, where *spacetime tells matter how to move*, we hypothesize that the geometry of CPT manifolds can similarly guide the dynamics of single cells in the transitions.

To extract dynamics information from snapshot sequencing data, Gioele La Manno and his collaborators proposed the RNA velocity method [4]. The RNA velocity method assumes simple dynamic balance between spliced RNA and unspliced RNA and estimates the time derivative of spliced RNA counts. This method has been further developed by several labs [5-7]. By integrating RNA velocity with Fisher metric, we can compute the information velocity of each individual cell in different coordinate systems (Fig. 6a) (Methods) [34-36], which quantifies the cell dynamics on the information manifolds.

**Figure 6.**
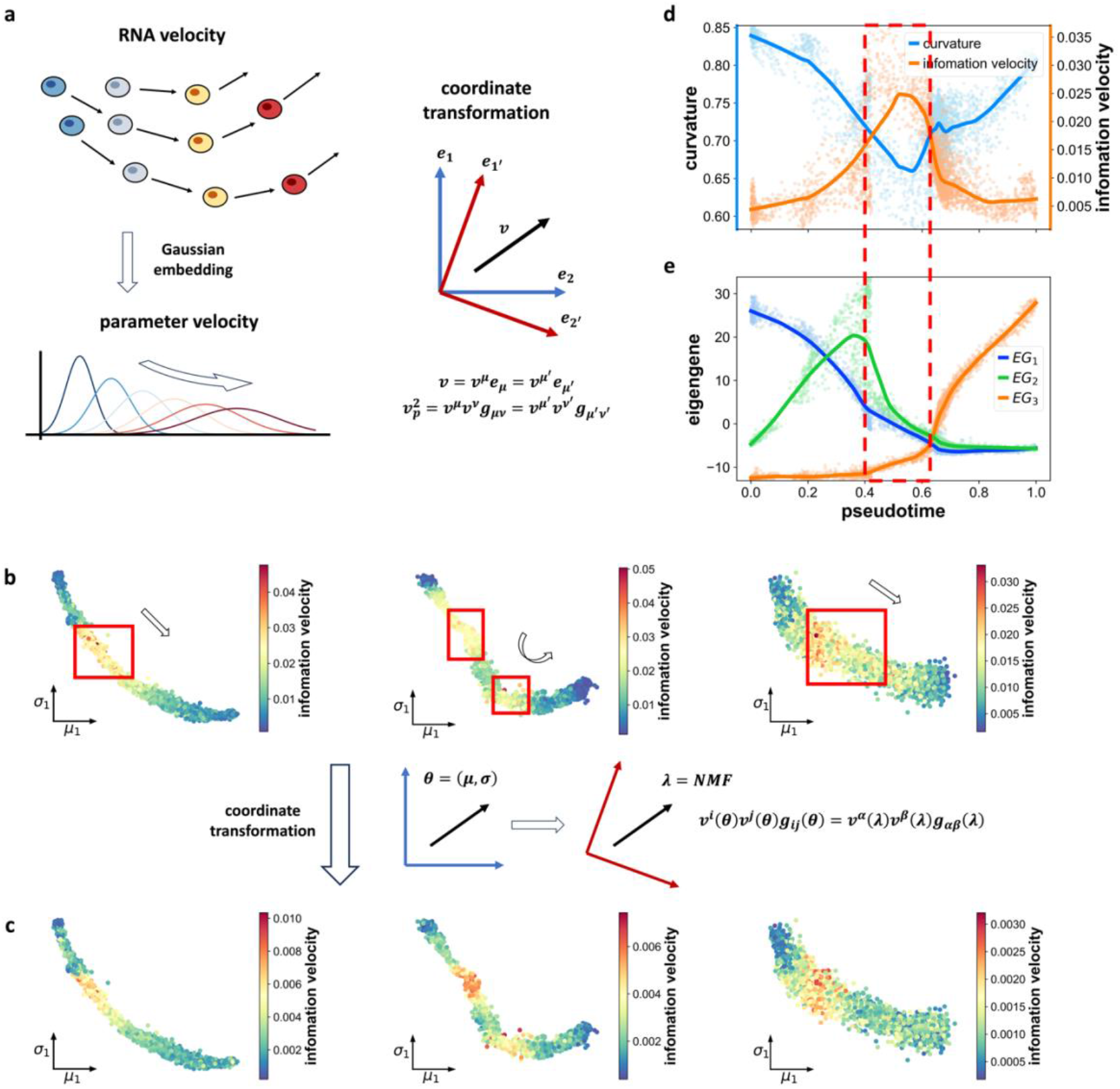
Information velocity analyses in different CPT processes with SCIM. (a) Flowchart of information velocity calculation using RNA velocity method. The RNA velocity is inferred from the dynamic equilibrium between unspliced mRNA and spliced mRNA. Information velocity is then calculated with the velocities and Fisher metrics. It can be calculated in different coordinates and is theoretically invariant under coordinate transformation. (*e*_1_, *e*_2_) and (*e*_1_′, *e*_2_′) denote different coordinates. *v* is the parameter velocity and *v*_*p*_ denotes the information velocity. (b) Variation of information velocity calculated within (*µ*, σ) coordinates in dentate gyrus neurogenesis (left), pancreatic endocrinogenesis (middle), and EMT of A549 cells induced by TGF-β (right). Cells with low CRC values mainly coincide with those exhibiting high information velocity values (boxes). (c) Variation of information velocity calculated within NMF coordinates in dentate gyrus neurogenesis (left), pancreatic endocrinogenesis (middle), and EMT of A549 cells induced by TGF-β (right). The similar distributions with that in panel (b) is consistent with the theoretical invariance of information velocity. (d) Association of low-CRC value and high information velocity in dentate gyrus neurogenesis (with respect to pseudo-time). The lines are fitted with LOWESS (Methods). (e) Association between manifold geometry and gene regulation networks (switch of eigengenes). Comparing with the minimum point of CRC and the peak of information velocity in panel (c), all three eigengenes show significant variations in slopes at this critical point (dashed box). The lines are fitted with LOWESS (Methods).

Analyses of various CPT processes revealed that cells in the low-CRC region display high information velocity values (Fig. 6b box), indicating a correlation between high information velocity and low CRC value (Fig. 6b). As the low-CRC region is around the saddle point, probability distributions on the opposite sides of the saddle point are distinct. Transitions between the two distributions are drastic, resulting in high information velocity values in this region. Conversely, low information velocity values in the basins indicates that the distribution shows little variation and cells are in stable states. These findings reflect that the geometry of underlying manifold guides the dynamics of single cells during CPT. Robustness checks using different parameters for CRC and information velocity calculations confirmed this conclusion (Fig. S10&S11). We also applied an alternative gene selection method named *DUBstepR* to preprocess all three datasets [62], yielding similar results of CRC and information velocity (Fig. S12) with that in Fig. 6d.

Additionally, we performed coordinate transformations and calculate information velocity in different coordinate systems. Using non-negative matrix factorization (NMF) for dimension reduction, we derived the Fisher metric in this new coordinate system. The velocities of each NMF dimension can be obtained by the linear combination of different velocity genes’ RNA velocities. We found that the distribution of information velocity is similar to that in the (*µ*, σ) coordinates (Fig. 6c and Fig. S13). From the definition of information velocity, it is evident that it is a value that is independent of coordinates. Our analyses confirm its theoretical invariance under coordinate transformation, although choice of coordinates can affect the absolute value of information velocity which is probably due to limitation of data and algorithm in practice. However, high information velocity value is not necessarily related to the high absolute value of RNA velocity. As shown by the distributions of norm of the RNA velocity in different datasets (Fig. S14), there is no direct correlation between them with that of the information velocity. When the space is Euclidean, the norm of RNA velocity vector is equivalent to the information velocity. However, this is typically not the case, as cell states usually reside on low-dimensional manifolds rather than in Euclidean space. Therefore, when analyzing cell state transitions on manifolds, it is crucial to consider the geometric properties of the manifold, such as metrics and curvature. Simply relying on the norm of RNA velocity or changes in gene expression vectors is insufficient to fully quantify the rate of cell state variations. By integrating the metric on the manifold, information velocity can more accurately capture the variation rate of cell states.

Eigengenes (EG) represent control modules in the gene regulation network, and switching between these modules is critical in cell state transition. Plotting eigengene values across the cells reveals a pattern that eigengenes form a relay in the dentate gyrus neurogenesis (Fig. 6e). Especially, the increase of EG3 is right after the decrease of EG1 and EG2. Furthermore, the low-CRC region or high information velocity region (Fig. 6d) coincides with the transitory region between eigengenes (Fig. 6e), suggesting the interaction between eigengenes or gene modules probably determine the geometry of CPT information manifolds. The dynamic pattens of CPT emerge from the complex interactions between genes. Specifically, the saddle region, characterized by low-CRC values and high information velocity, probably involves more gene interactions even global reprogramming of gene profiles [53, 63].

### Characterizing branching points with Fisher information

In this work, we primarily analyzed the transition of cells from one phenotype to another. However, stem cells often differentiate into multiple cell types, leading to complex critical transitions and manifolds. Identifying these branch points is crucial for lineage reconstruction in embryo development. We also tested our method on datasets with different branching structures. First, we applied it to a toy dataset with two branches (Fig. 7a). The CRC value at the branch points was low (Fig. 7b). As low curvature is associated with increased Wasserstein distance (Fig. S7), the stiff dimensionality of Gaussian embedding, which influences cell state, should correspondingly enhance its counterpart in Wasserstein distance. Additionally, we observed an increased negative correlation between Fisher information in the local neighborhood across different stiff dimensionalities. This negative correlation suggests a switch in stiff directions within the parameter space, whereby some dimensionalities become stiffer while others become less so. For each dimensionality, we calculate its total negative correlation value (as an absolute) with all other dimensionalities, referred to as negative correlation strength. When both its counterpart in Wasserstein distance and negative correlation strength demonstrate high values within a single cell, it is classified as a varying stiff dimensionality for that cell (Methods). The ratio of varying stiff dimensionality to latent dimensionality *L*, peaks around branch points (Fig. 7c). This varying stiff dimensionality ratio highlights the regulatory variation surrounding branch points.

**Figure 7.**
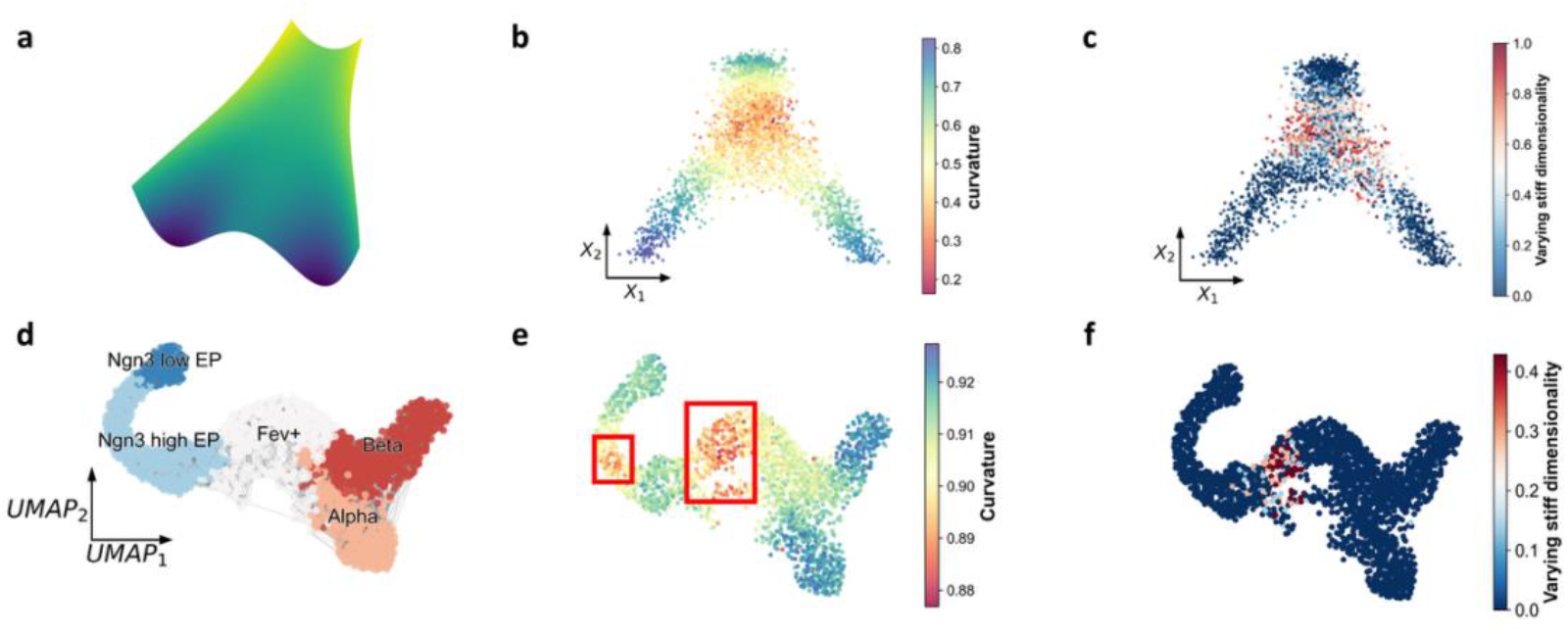
Application of SCIM on data with multiple branching structures. (a) Synthetic data generated through simulations on landscapes with multiple branching structures. (b) CRC distributions across synthetic data points. (c) Ratio of varying stiff dimensionality number to embedding dimensionality *L* in each cell. Cells near the branch points show high ratios, indicating increased number of varying stiff parameters at these points. (d) Cell graph of endocrinogenesis data including both α and β cells. (e) CRC distribution reveals two transition regions in this process. The CRC analysis identifies two distinct transition regions, consistent with previous analyses on single branches. (f) Ratio of number of varying stiff dimensionality to embedding dimensionality *L* in each cell. Similar to the synthetic data, cells in the transition region around the branch point exhibit high ratios, while cells in the other transition region only show slight increase.

During development, pancreatic endocrine cells differentiate into various branches, including glucagon-producing α-cells and β-cells (Fig. 7d). We observed that the distribution of CRC is essentially the same as in Fig. 5d when the dataset includes only the α-cell branch. The two low-CRC regions correspond to transitional states (Fig. 7e). However, CRC alone cannot differentiate between the two transitional states, as the latter represents a branch point. We calculated each dimensionality in Wasserstein distance between neighboring cells (Fig. S15 a) and negative correlation strength in all cells (Fig. S15 b). We analyzed the ratio of varying stiff dimensionality and found that the latter transitional region exhibited a significant increase in this ratio (Fig. 7f). We also observe slight increase in the first transition region when using a larger latent dimensionality in embedding (Fig. S15 c). This difference indicates transition at branch point involves more varying stiff parameters since it connects more cell states. The result that varying stiff dimensionality peaks around branch points is robust under different settings of parameters and random subsampling (Fig. S16).

While information velocity cannot directly identify branch points, we projected the information velocity onto the coordinates of different eigengenes corresponding to different branches (Fig. S17). This projection helps distinguish the branch points by highlighting changes in the underlying gene regulatory networks.

## Discussion

The exploration of underlying manifolds of CPT has recently emerged as a novel research field. In this study, we have developed a method based on information geometry to investigate the intrinsic geometric properties of the CPT manifolds. Our approach, the Single-Cell Information Manifold (SCIM) method, diverges from traditional manifold learning methods that focus solely on low-dimensional visualization. Instead, SCIM examines the CPT manifold through its intrinsic geometry, independent of visualization techniques. It transforms the single cell gene vectors into multivariate Gaussian distributions, allowing for the computation of Fisher metric and CRC of information manifold in probability space. We discover that the geometric properties of manifold, such as CRC, correlates with dynamical properties like information velocity in CPT. Moreover, the CRC distributions are consistent across different Gaussian embeddings and information velocity empirically exhibits invariance under coordinate transformations.

When calculating the Fisher metric, constructing a cell graph from experimental data is necessary. However, sampling biases can affect the accuracy of this step. For instance, the local cell density variations probably alter the neighbor diversity in kNN graph, potentially impacting the computation of the Fisher metric. Although we do not find direct correlation between CRC and local cell density, the sampling bias should be taken into consideration, particularly when dealing with time-course datasets like the EMT dataset of A549 cells. Moreover, as experimental techniques constrain sampling, extrapolation the analyses into the regions where there are no sampled cells is not feasible. Our focus is on leveraging local neighborhood information to investigate the properties of single cells. The computational complexity related to data scale primarily stems from the construction of kNN graph, which has a time complexity of O(n^2^m),where n is the number of cells and m is the number of genes. In datasets used as examples, the maximum number of cells and genes are on the order of 10^4^ and 10^3^ respectively, which can be managed with typical computing resources. For datasets with significantly larger numbers of cells, a potential strategy is to initially apply k-means clustering to partition the data into several thousand or even tens of thousands of cell subpopulations. Subsequent SCIM analyses can then be performed on the centroids of these subpopulations. This approach is suitable when the analytical focus is on regions rather than individual cells and does not introduce significant distortion. A similar method has been employed in *Mutrans* for managing large-scale datasets [64].

The intrinsic geometry of the CPT manifold is related to the intrinsic dimensionality, the estimation of which is still an open question [65]. An accurate intrinsic dimension can increase the training efficiency and accuracy of neural networks, while an incomplete coordinate system may distort the calculation of the Fisher metric. We vary the number of latent dimensions to ensure that the transformation is robust (Fig. S10). But the methodology for estimation of intrinsic dimension and its reliability will be further explored.

Duluxan Sritharan and colleagues proposed a method to calculate Riemann curvature with extrinsic differential geometry in single cell RNA sequencing data [66]. Different from the extrinsic geometry method, SCIM is rooted in intrinsic geometry which is invariant under coordinate transformation. It allows us to study the system by transforming into different biological meaningful coordinates, such as eigengenes.

Another prevalent technique for embedding high-dimensional data with probability distribution is diffusion map [25], which employs kernel based method for constructing Markov transition matrix between neighboring data points. PHATE further develop this approach with information-theoretic potential distance for improved visualization of high-dimensional biological data [19]. Oluwadamilola Fasina and colleagues developed neural FIM with Markovian diffusion operator based on PHATE for computing Fisher information metric [67]. While neural FIM represents data points with diffusion probability distributions and defines Fisher metric on the original feature space, SCIM captures the uncertainty of individual cells with the transformed Gaussian distributions and learns Fisher metric in various coordinate systems through transformation.

Nowadays, RNA velocity has become a potent tool for learning the dynamics of CPT. However, it relies on some strong assumptions and suffers from some flaws when applied to different datasets [68]. Considering that the geometry of the CPT manifold is related to the cells’ dynamics, this method can be further refined to regularize the inferred dynamics with geometric properties. For instance, the fact that information length computed with information velocity and with local Fisher metric should be consistent can be used as constrain on inferring RNA velocity. Additionally, RNA velocity relies on the counts of unspliced and spliced RNA for its inference. When this information is unavailable, alternative approaches, such as cell velocity based on optimal transport, can be employed to calculate information velocity [31].

Study of critical points in complex systems is still an active research field [55, 56, 69, 70]. Information geometry, along with Fisher information, offers a powerful means to characterize phase transitions and can be readily adapted to non-equilibrium systems because it does not require defining Hamiltonian or ensemble function [71, 72]. The peaks observed in the non-diagonal elements of the Fisher information matrix are similar with the divergence of order parameter in phase transition, which indicates CPT probably share similar characteristics with phase transition (Fig. 3d and f) [71]. Therefore, SCIM not only offers a framework for analyzing single cell transitions, but may also reveal underlying principles of transitions in a broader spectrum of complex systems [72, 73]. While SCIM uncovers certain properties of branch points, the calculation of orthogonal stiff parameters and their corresponding biological functions requires further exploration.

The manifolds may evolve due to intricate interactions among cells or external signals in complex biological processes. In all the analyses in this work, the parameter systems do not explicitly contain time. However, some parameters implicitly contain information about the direction of time. For example, the EMT dataset is collected from different time points and clearly shows that *μ*_1_ is correlated with time (Fig. 4h). And the parameters that control bifurcation like X_2_ in Fig.7 can be also time dependent [53, 55]. Currently, we use all the data to learn the geometry only in spatial coordinates. The exploration of pseudo-Riemannian geometry, which incorporates both time and space, will be pursued in future studies. Quantitative descriptions and analyses of the geometry of evolving manifolds can help us characterize different stages in complex developmental processes.

## Figures

## Methods

### 1 Gaussian embedding of single cell RNA sequencing

Single cell RNA sequencing is a powerful technique that measures abundance of mRNA in individual cells, enabling a detailed understanding of cell heterogeneity. Each cell is characterized as a point in a high-dimensional gene space. We construct a k-nearest neighbor (kNN) graph by computing pair-wise Euclidean distances between cells in this space (Fig. 1a). The number of neighbors used in kNN cell graph is denoted with *k*_*nei*_ .

To represent each cell as a probability distribution, we employ the Gaussian embedding method, a technique previously applied in natural language processing [33]. The SCIM framework utilizes neural networks to learn the Gaussian embedding of each cell based on its high-dimensional gene vectors and its proximity to neighboring cells in the gene space *R*^*D*^ ( *D* is the original high-dimensional gene expression space) (Fig. 1b) [32]. Each cell is transformed into a multivariate Gaussian distribution in an *L*-dimensional space *R*^*L*^ (Fig. 1c), where *L* is significantly lower than *D*. For simplicity, the covariance matrices of transformed Gaussian distributions are diagonal.

In this study, the neural network is trained with an energy-based learning method. The energy-based model is a method of capturing dependencies between variables through mapping each configuration of the variables to a scalar energy. And the loss function is used to evaluate the qualities of different energy functions. During training, triplets (*i, j*_*k*_, *j*_*l*_ ) as illustrated in Fig. 1b are sampled uniformly from the cell graph. The individual cells’ gene vectors are used as inputs in the neural network. The loss function is defined as

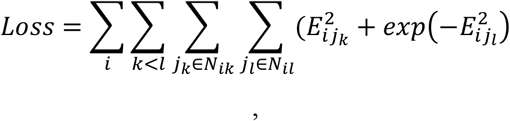

where *N*_*ik*_ represents the set of nodes *j*_*k*_ that the length of shortest paths on the graph between node *i* and node *j*_*k*_ equal *k*. The maximal *k* considered is denoted by*k*_*hop*_. The energy *E*_*ij*_ represents the distance between nodes *i* and *j*. By minimizing this loss, the 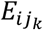 in each term is minimized as much as possible while 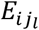 is maximized.

In the original work, the energy function is Kullback-Leibler divergence *E*_*ij*_ *= D*_*KL*_ (*P*_*i*_||*P*_*j*_) [32]. In our approach, we use the Fisher distance as the energy function,

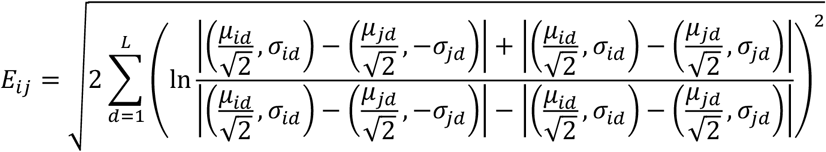

where (*µ*_*id*_, σ_*id*_)represents the coordinate of the *d*_*th*_ component of the *i*_*th*_ Gaussian distribution. In this way the geometric structure of data points represented by the cell graph is preserved into the embedded Gaussian distributions. Once the neural network is trained, it takes single cell’s gene vector as input and transforms this vector into a Gaussian distribution in application.

Typically, we take *k*_*nei*_ *=* 10, *L =* 10, *k*_*hop*_ *=* 2 for experimental data. Guidance on parameter selection can be found in the [Readme.md](https://github.com/wwklab/SCIM/tree/main) file on Github. We also test other parameters (Fig. S9&S10), and the result is robust to the change of the parameters.

In contrast, the loss function of VAE is *loss =* −*E*_*z*_[*p*(*x*|*z*)] + β*KL*(*pθ*(*z*|*x*)||*q*(*z*)), where *x* represents the input data, *z* is the variable of the embedded distribution and β is the weight of KL-divergence. As can be seen, the second term pushes the embedded distribution *pθ*(*z*|*x*) to a given prior *q*(*z*), so keeping geometric structures in latent embedding space is not the goal of VAE embedding.

### 2 Riemannian manifold with Fisher metric

The single cell RNA sequencing data typically forms a manifold structure, which means the data points are concentrated to a hypersurface in the gene space, and the manifold typically is low-dimensional. Manifolds can be quantified by adding a metric tensor, which specifies the infinitesimal distances on it. A manifold equipped with a positive-definite metric tensor is called a Riemannian manifold. However, in many cases, there is no meaningful way to define the metric. By representing single cells as Gaussian distributions, there is a natural way to define the metric. Using theories from information geometry, we calculate the Fisher information of each cell, which acts as the Riemannian metric in the probability distribution space. The Fisher information matrix is defined as

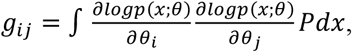

where *θ* are the parameters of the distribution and *g*_*ij*_ represents an element of the Fisher information matrix. Here *θ* is the general symbol representing parameters. For 1-D Gaussian distribution, *θ =* (*µ*, σ) and the Fisher information of *µ*and σare 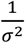 and 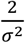 respectively [26, 37]. And the Fisher information matrix *G* of a *L*-dimensional diagonal Gaussian distribution is

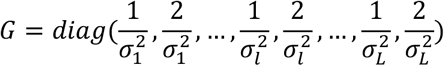

where σ_*l*_ represents the standard deviation of *l*^*th*^ dimensionality of Gaussian distribution. For clearance, each cell has a Fisher information matrix as each single cell *X* is transformed into a Gaussian distribution N(***µ*, σ**).

Defining connections such as Levi-Civita connection, which is fundamental to curvature in this space, remains a complex challenge when working with experimental data. Therefore, we employ an alternative method proposed by Yann Ollivier, which estimates the curvature on a graph by assigning a probability measure to each node, and then calculates the coarse Ricci curvature (CRC) of each edge using the following equation [39]:

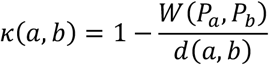

where *W*(*P*_*a*_, *P*_*b*_) represents the Wasserstein distance between the probability measures of node *a* and node *b* and *d*(*a, b*) is the length of shortest path between node *a* and node *b* [38]. The Wasserstein distance measures the minimum cost of transportation required when transforming one probability distribution to another. Ricci curvature, on the other hand, controls the overlap of two balls or the overlap between distributions, and is directly related to the Wasserstein distance [39].

By representing each single cell as a multivariate Gaussian distribution with mean ***µ*** and diagonal covariance **Σ**, we can directly calculate the Wasserstein distance between neighboring cells:

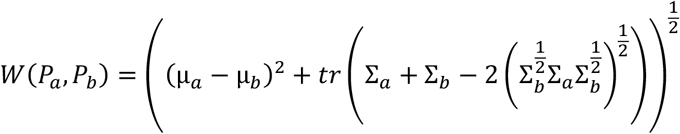

and *d*(*a, b*) is the Euclidean distance between neighboring cells in gene space. Then we can calculate CRC of each edge, and the CRC of each cell is then determined by the average of CRC across all its connecting edges 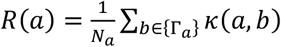 . Here *{*Γ_*a*_ } represent all the neighboring cells of cell *a* and *N*_*a*_ is the number of neighbors of cell *a*. Here we used 2-Wasserstein distance (also referred to as the *L*^2^ transportation distance in [39]) in calculating CRC as the multi-variate Gaussian distribution has closed form of 2-Wasserstein distance which is easy for calculation. As we have used the transformed Gaussian embedding in calculation, the CRC value we obtained is an approximation of the original definition by Yann Ollivier.

### 3 Test data generation

In our research, we use a series of toy models to generate data for validation of SCIM approach. Specifically, we employ four different models:

#### (1) Saddle surface

Data points are randomly sampled from a saddle surface, a two-dimensional manifold embedding in a three-dimensional space, represented by the equation *z = k*(*x*^2^ − *y*^2^). Here we take *k =* 0.*5* . To these points, we add seven additional dimensions of Gaussian distributed noise, which introduce minor variances in the data compared to the dominant first three dimensions. Despite the introduction of noise, the resulting ten-dimensional data points remain to be confined on the 2D saddle surface.

### (2) Ellipsoid surface

We randomly sampled data points on an ellipsoid surface, where the original data are three-dimensional and defined by the equation:

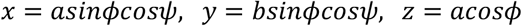

(*Φ*, ψ)are angles and (*a, b*) are the length of the semi-axes (*a =* 1, *b =* 4).

As with the saddle surface model, seven dimensions of noise were added to the original three-dimensional data.

#### (3) Four-well potential landscape

Data points were obtained from a four-well potential surface described by the function *U = a*(*kx*^4^ − *x*^2^ + *ky*^4^ − *y*^2^) (where *U* is the potential), generated through trajectory simulations using the Langevin equation. Here we take *a =* 0.*5, k =* 0.05 . Subsequent to the simulation, the trajectory data are added with seven dimensions of noise, in a manner consistent with the previous models.

#### (4) Simulated differentiation process with double branches

Data points were sampled with trajectory simulation on the double-branch potential surface described by the function *U = k*_1_*x*^4^ + *k*_2_*yx*^2^ + *k*_3_*y* (where *U* is the potential). The simulations were terminated when the range of *y* exceeds 35.

Here we use *k*_1_ *=* 0.05, *k*_2_ *=* −0.06, *k*_3_ *=* −1. The trajectory data were also added with seven dimensions of noise.

By applying SCIM to these toy models, we can assess its feasibility and effectiveness in exploring the manifolds of CPT. These models allow us to evaluate the performance of SCIM in different scenarios and gain insights into its advantages and limitations.

### 4 Comparison of local metric preservation capability

To evaluate the preservation of metric in original space, we sampled points on the ellipsoid surface with equal major and minor axes (a=b=1) i.e. a sphere surface. We then sampled multiple semicircles on this surface, which represent geodesic lines on the sphere.

We first calculate the length of these geodesic lines (*L*_*x*_) on the kNN graph using shortest path method in *Networkx* (Dijkstra algorithm). The weights assigned between neighboring nodes correspond to the pairwise Euclidean distances (*d*_*x*_) and the lengths of these semi-circles approximate π, although the true values are subject to perturbation by seven-dimensional noise. Subsequently, we calculated the length of semi-circles (*L*_*F*_) in the embedding spaces. For Gaussian embedding, the length is calculated with pairwise Fisher distances(*d*_*F*_) . For other manifold learning methods, since they lack defined metrics, we use the pairwise Euclidean distances (*d*_*E*_) within the embedding space to compute the length of semi-circles(*L*_*F*_) . By comparing the distribution of *L*_*F*_/*L*_*X*_ with that of *L*_*E*_/*L*_*X*_, we assess the effectiveness of the Gaussian embedding in maintaining the integrity of the original local metric. And the coefficient of variation (CV) of the ratio distribution provides a quantitative measure of this comparison.

### 5 Coordinate transformation of Fisher matrix

We use a neural network to learn the mapping function between vector pairs. Specifically, we train a simple multi-layer perceptron neural network (two hidden layers) with *PyTorch*, where the input is the user-defined coordinates *λ* and the output is the (*µ*, σ) (represented with *θ* ) of the Gaussian distribution. The derivative of this mapping function *θ = f*(λ) can be estimated with *autograd* in *PyTorch* numerically. This allows us to compute the transformation between (*µ*, σ) and other coordinates. With the following equation

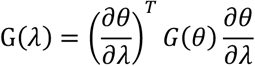

Fisher metric G(λ) in the coordinates *λ*with biological meaning such as eigengenes and principal components can be computed, thereby facilitating the quantitative analysis of CPT.

The Fisher information matrix in the (*µ*, σ) space is diagonal. However, this is not necessarily the case in other coordinate systems.

### 6 Identification of stiff and sloppy genes

Fisher information can also be employed to measure the stiffness and sloppiness of different parameters. The probability distribution is highly sensitive to changes in stiff parameters, whereas sloppy parameters have minimal impact on the distribution. To calculate the Fisher information of different genes, we need to perform coordinate transformation from (*µ*, σ) space to gene space:

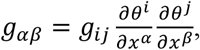

where *θ =* (*µ*, σ) are the parameters, and *x* represent genes (here not the variables of Gaussian distribution). And Einstein summation convention is used in this equation. Instead of directly performing this transformation, we first transform to the PCA space and then to the gene space:

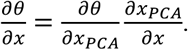

Particularly we focused on the Fisher information of each gene. The Fisher information of gene 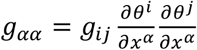 (Einstein summation convention is applied).

### 7 Information velocity and Information length

Information length is defined as the accumulated variation of information along a path in the manifold and is given by this equation,

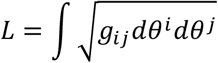

where 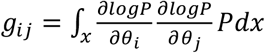 is the element of the fisher information matrix G, and *P = P*(*x*, t) is a distribution function which is Gaussian embedding in current context (Einstein summation convention is applied). In current context, information length represents the accumulated variation of cell state. This concept is widely used in analyzing thermodynamics and attractor structure [34, 35].

Information length can also be calculated with integration of information velocity along a path.

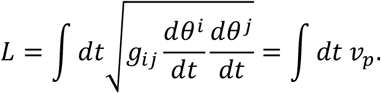

By applying the chain rule twice, the information velocity *v*_*p*_ is defined and calculated with the following equation:

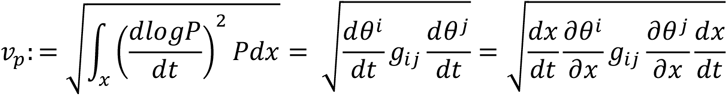

Where 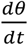 represents the velocity of the parameter and *θ* is (*µ*, σ) in Gaussian distribution, 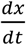 represents RNA velocity. Information velocity measures the variation rate of the probability density function and reflects the speed of information variation along a path [34-36].We calculate the information velocity of each cell using RNA velocity. RNA velocity reflects the direction of single cell along the path of CPT in the gene expression space (*X*), thus information velocity of single cell represents the speed of information variation along the transition path of CPT in probability distribution space. With RNA velocity 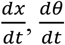 can be expressed via chain rule 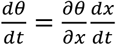 . The 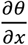 is calculated using *Pytorch autograd* function numerically. For the eigen-gene or PCA coordinates (λ), the velocity is defined as 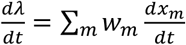, where *w*_*m*_ is the weight of velocity gene *x*_*m*_ in the corresponding coordinates.

### 8 Eigengene analysis of regulatory mechanisms of CPT manifolds

To investigate the underlying regulatory mechanism of CPT manifolds and their characteristics, we compute the co-expression profile of genes based on their correlations. Utilizing the gene-gene correlation as a measure of distance, we conducted hierarchical clustering and separating the genes into distinct modules. The eigengene (EG) of each module is defined as its first principal component (PC1) value *EG = W*_*m*_*X*_*m*_ [75, 76], representing the collective behaviors of genes, where *W*_*m*_ is weight vector of gene module *X*_*m*_. These eigengenes reveals the regulation of underlying gene networks and are treated as parameters that modulate the probability distributions in our analysis.

### 9 Identification of varying stiff dimensionality

Considering the diagonal Fisher information matrix of Gaussian embedding, the counterpart of each dimensionality of Gaussian embedding in Wasserstein distance between neighboring cell can be easily calculated.

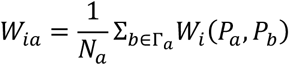

Where i, a, *N*_*a*_, Γ_*a*_and *P*_*a*_represent the index of direction, index of cell, number of neighboring cells of cell *a*, neighboring cells of cell *a* and Gaussian embedding of cell *a*.

For each dimensionality of Gaussian embedding, the Fisher information for μ and σ are 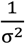 and 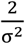 . The negative correlation strength 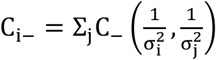 of dimensionality *i* is defined as the total absolute values of its negative correlations with other dimensionalities.

Then we use Otsu algorithm to label cells with high values of *W*_*ia*_ and *C*_*i*−_ [77]. A cell is labeled as 1 if both its *W*_*ia*_ and *C*_*i*−_ are above Otsu threshold values respectively, dimensionality *i* is defined as varying stiff dimensionality in this cell. The ratio of number of varying stiff dimensionality to L is calculated in each cell.

### 10 Clustering method and evaluation

For cell type clustering, we use the diagonal element of the FIM of each cell as a vector to represent the cell and use the vectors’ cosine distance to perform agglomerative hierarchical clustering to obtain the clustering results. Agglomerative hierarchical clustering is a bottom-up clustering algorithm. It begins by treating each data point as an individual cluster. The algorithm then iteratively merges the two most similar clusters, continuing this process until all data points are merged into a single large cluster or a predefined stopping condition is met, such as specifying the desired number of clusters [78].

In addition to testing on the dentate gyrus and pancreatic endocrinogenesis datasets, we also evaluated these approaches on a zebrafish dataset with cell type labeling [79].

Three metrics were used for evaluation of the clustering performance. Rand index (RI) evaluates the match between the clustering results and the true label by statistically analyzing the distribution consistency of sample pairs. Mutual Information (MI) measures the degree of information sharing between the clustering results and the true label based on information entropy. ARI and AMI are the results of their correction based on the expected value of the random distribution cluster label assumption. F1-score is the harmonic mean of recall and precision. Larger is better for these clustering performance metrices. Here are formulas of these comparing performance metrics:

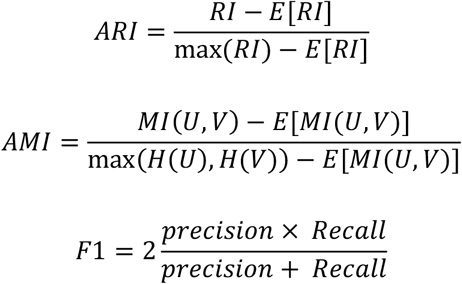

Where *U, V* means ground truth label and clustering label, *H* means entropy and *E*[] means expectation of the random distribution.

### 11 Go enrichment

We used *gseapy* package to implement the GO Enrichment Analysis [61]. Our gene selection thresholds were set at p-value < 0.05 and FDR < 0.05, using the KEGG_2019_Mouse database as a reference. Significant terms were defined as GO terms with an FDR-adjusted q-value < 0.05. Here, we presented the top six terms ranked by combined score.

### 12 Density estimation

To estimate local cell density, we employ a method using k-nearest neighbor and Gaussian kernel [80]. The local density is defined with 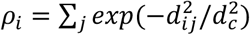, where *d*_*ij*_ is the distance between cell *i* to its neighbor cell *j* and *d*_*c*_ is a cutoff distance. We set *d*_*c*_ as the mean distance between all neighbor cell pairs. Given the experiment data is often noisy, we incorporate a smoothing step to refine our analyses. For a scaler function *f* like curvature and information velocity defined on single cell, its value is smoothed by the following equation 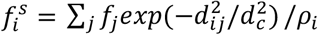 .

### 13 Locally Weighted Scatterplot Smoothing (LOWESS)

For better visualizing the variation of CRC or information velocity, we use LOWESS in *Scipy* to model the relation between the CRC or information velocity and pseudo-time. LOWESS is a non-parametric regression method that employs local linear estimation for smoothing scatter plot, providing a better view of the underlying trends in data.

### 14 Pseudo-time analysis

The pseudo-time in all the single cell RNA sequencing datasets are determined with Palantir [81]. The Palantir method employs a probabilistic method for characterizing cell fate. With Palantir, we establish a pseudo-time ordering for cells within the single cell RNA sequencing dataset. The dataset for EMT of A549 cells includes samples that treated with different durations with TGF-b. Unlike actual experiment time, the pseudo-time uncovers the heterogeneity of single cell dynamics on a more granular temporal scale.

### 15 Dataset preprocessing

The single cell RNA sequencing datasets of dentate gyrus neurogenesis, pancreatic endocrinogenesis, and EMT of A549 treated by TGF-b were obtained from the GEO website with accession number GSE95753 [42], GSE132188 [45], GSE121861 [46] respectively.

To concentrate on the process of CPT within each dataset and minimize the influence of genes that are not associated with transition, we use two methods to select the genes exhibiting significant variations during CPT. The first one is selecting genes that display switch-like behaviors by using approach described in [63]. All results, except for Fig. S12, were obtained using data processed with this method. The second method is DUBStepR, which is based on gene correlation analysis and helps filter out those genes that are not cell-type-specific (Fig. S12) [62]. We run 30 regression steps to select seed genes which can best explain the gene-gene correlation, iteratively add the most correlated gene to this seed gene set. For the dentate gyrus data, we used 600 DUBStepR-identified genes and 678 switch-like genes, with 368 genes overlapping. In the pancreatic endocrinogenesis dataset, there were 757 DUBStepR genes and 470 switch-like genes, with 348 shared genes. For the EMT of A549 data, 500 DUBStepR genes and 510 switch-like genes were identified, with a total of 242 shared genes. These results demonstrate the robustness of SCIM across different feature selection methods.

## Code availability

The scripts are available on GitHub at https://github.com/wwklab/SCIM.

The authors declare no conflict of interest.

## Acknowledgement

W.W. is supported by National Natural Science Foundation of China Grants No.12247104 and National Innovation Institute of Defense Technology Grant No. 22TQ0904ZT01025. L.Z. is supported by the National Key Research and Development Program of China 2024YFA0919500 and National Natural Science Foundation of China (No. 12225102, T2321001, 12288101, and 12426653). J.Y. is supported by Beijing Natural Science Foundation No. QY24006. We also thank Dr. Li Li (Institute of Theoretical Physics, Chinese Academy of Sciences) for insightful discussion on Riemannian geometry.

