## supplement for "Geometric Quantification of Cell Phenotype Transition Manifolds with Information Geometry"

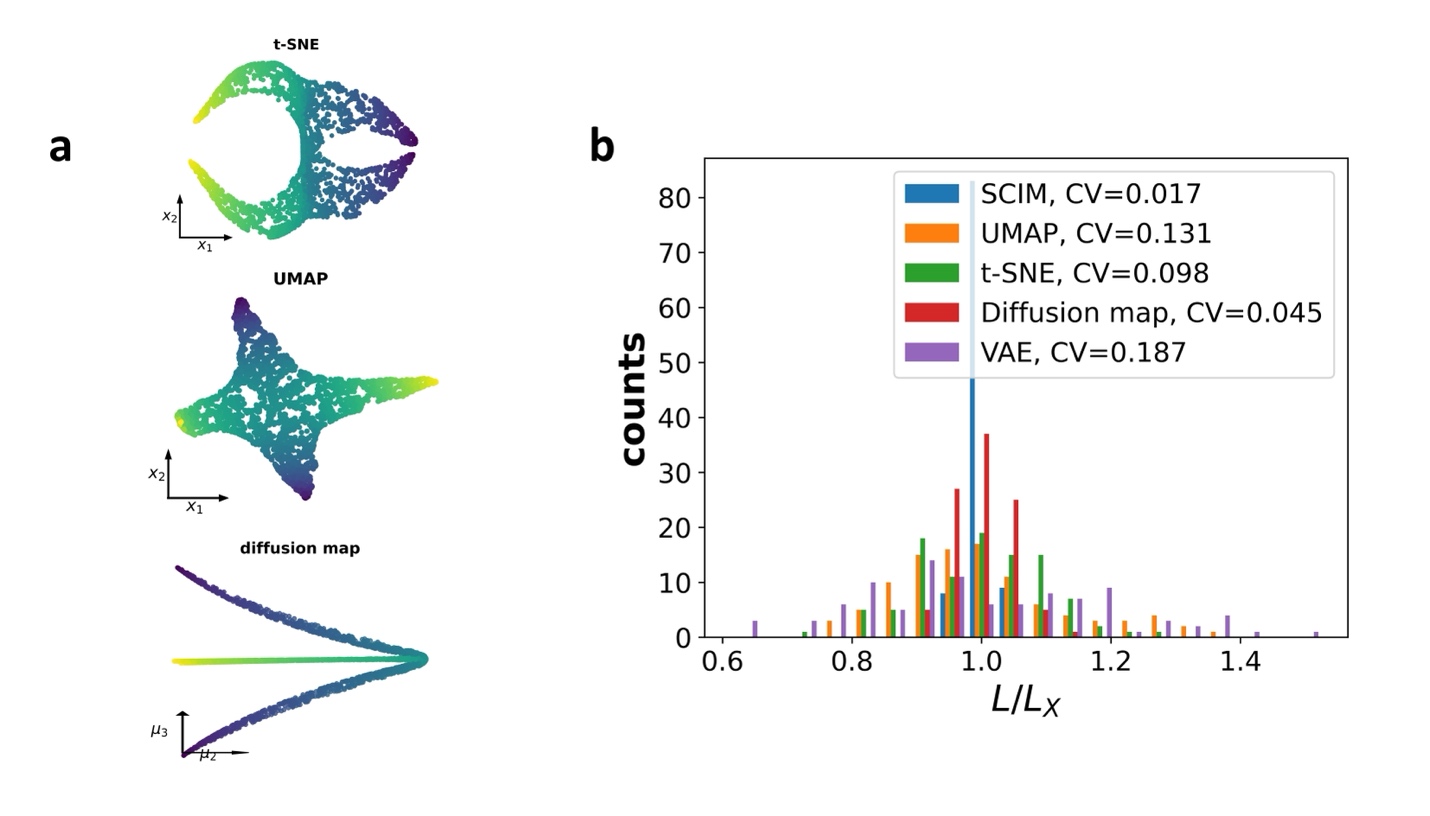
**Figure S1** Comparison of different embedding methods on toy datasets.

(a) Embeddings of saddle surface with t-SNE (top),UMAP (middle) and diffusion map (bottom).
(b) Comparisons between distributions of semi-circles length ratios before (*L_X_*) and after (*L*) embedding with SCIM, UMAP, t-SNE, diffusion map and VAE. The SCIM-based distribution is characterized by less dispersion (minimum coefficient of variation (CV)).

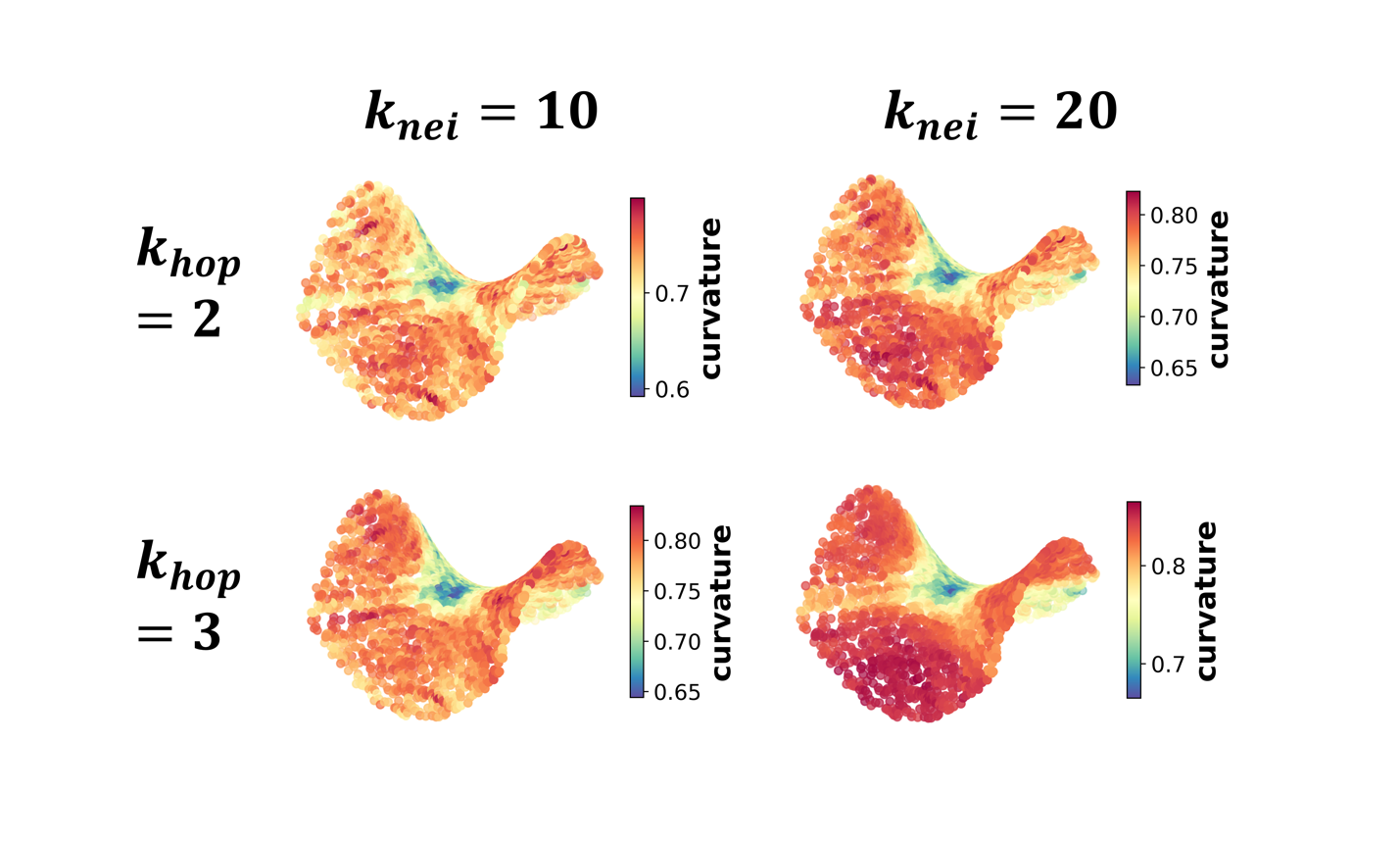
**Figure S2** CRC values of saddle surface are computed using different values of $k_{nei}$ and $k_{hop}$.

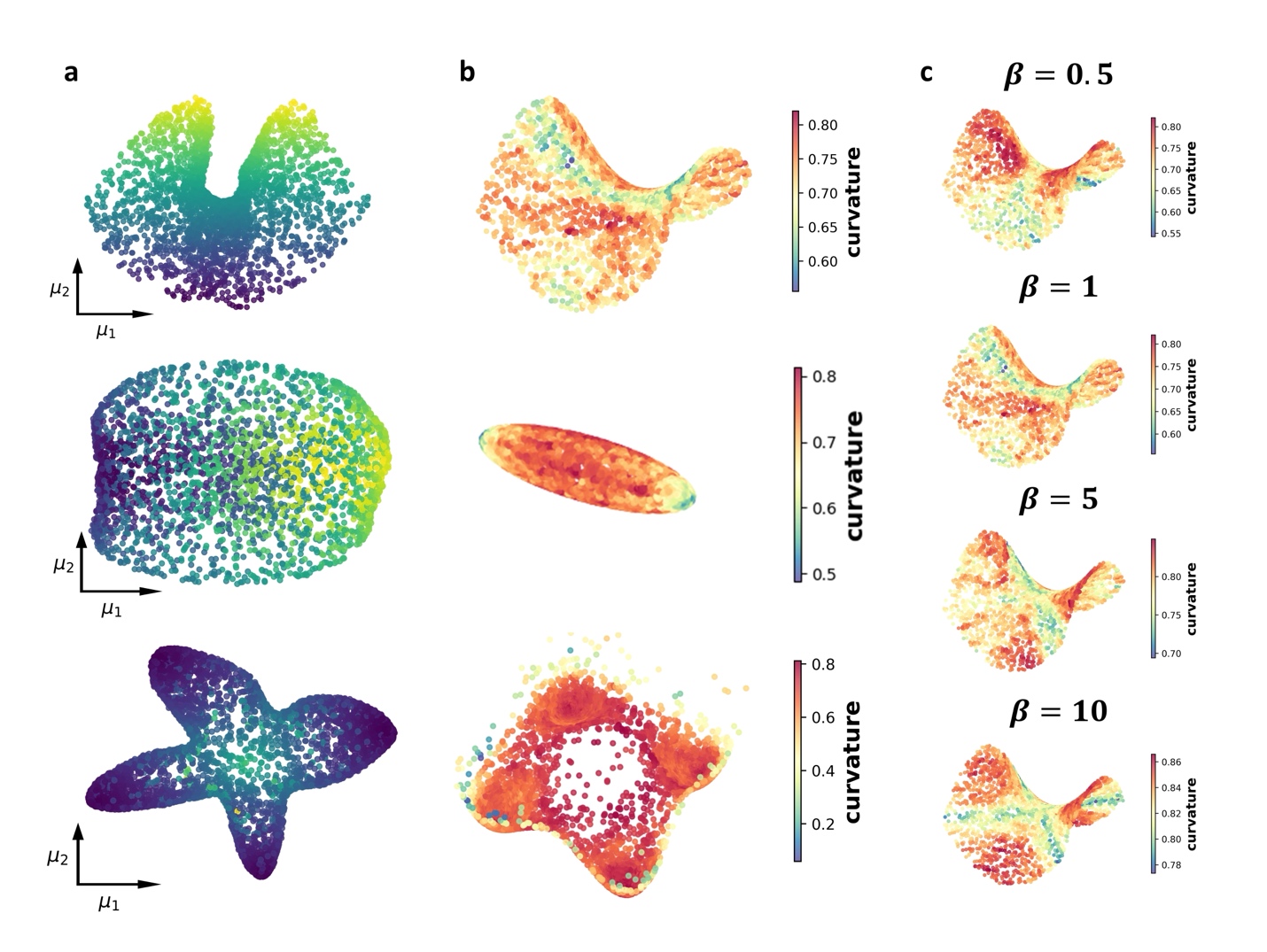

**Figure S3** Application of VAE on the toy dataset.

(a) Embedding of toy datasets with VAE: saddle surface (top), ellipsoid surface (middle), and four-well potential landscape (bottom).
(b) CRC distributions computed based on the embedding of VAE in saddle surface (top), ellipsoid surface (middle) and four-well potential (bottom). Color represents the value of CRC.
(c) CRC distributions of saddle surface computed based on the embedding of VAE when varying the relative weight (b) of KL-divergence in loss function.

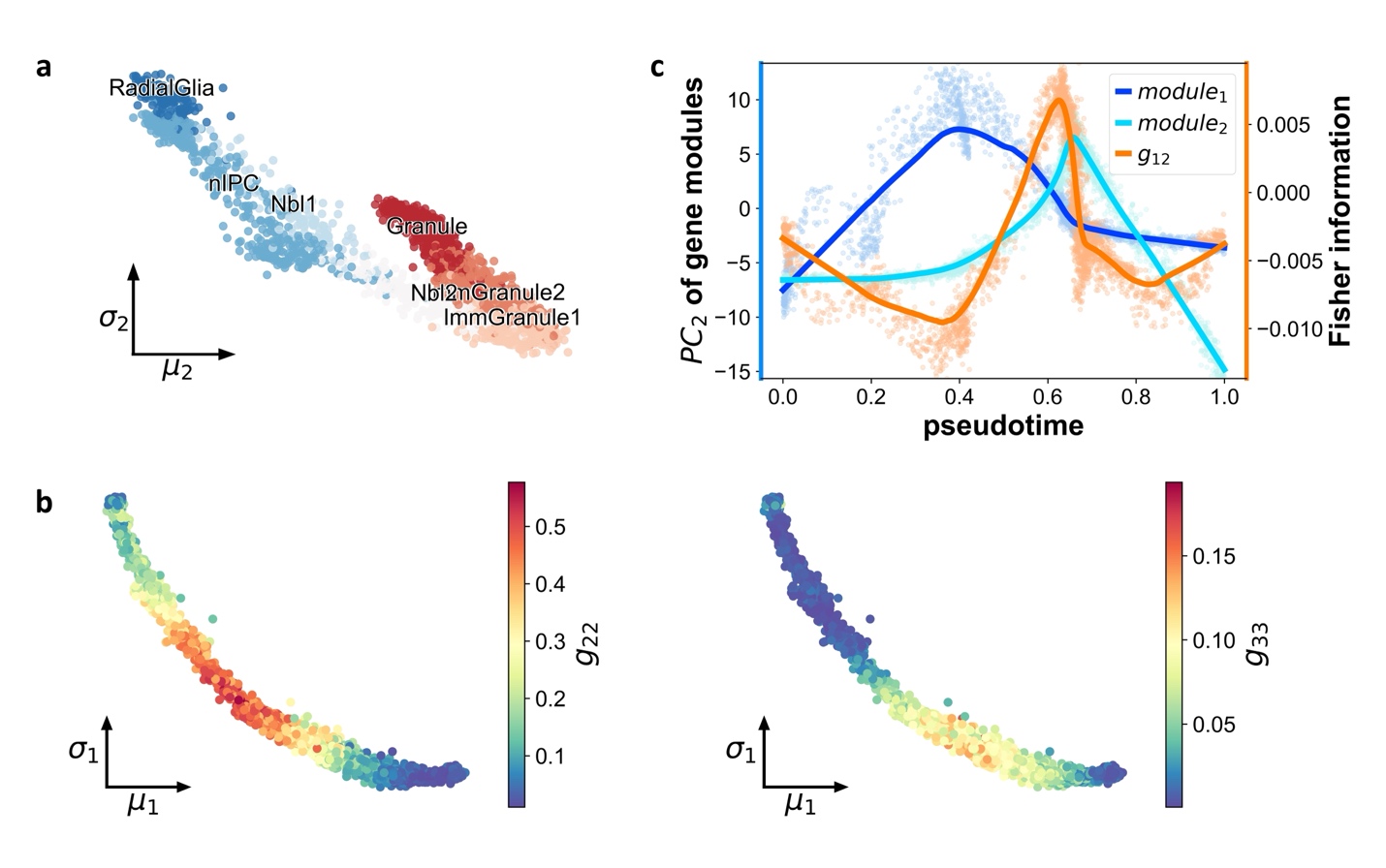

**Figure S4**

(a) Projection of dentate gyrus neurogenesis onto the plane of $(\sigma_{1},\sigma_{2})$ with SCIM.
(b) Variation of Fisher information of $\mu_{2}$ and $\mu_{3}$ (the second and third diagonal elements of Fisher matrix) in dentate gyrus neurogenesis. Color represents value of Fisher information.
(c) Variation of Fisher information of second principal component (PC2) of two gene modules (analogy with eigengene) and the corresponding values of PC2 (with respect to pseudo-time). The lines are fitted with LOWESS (Methods).

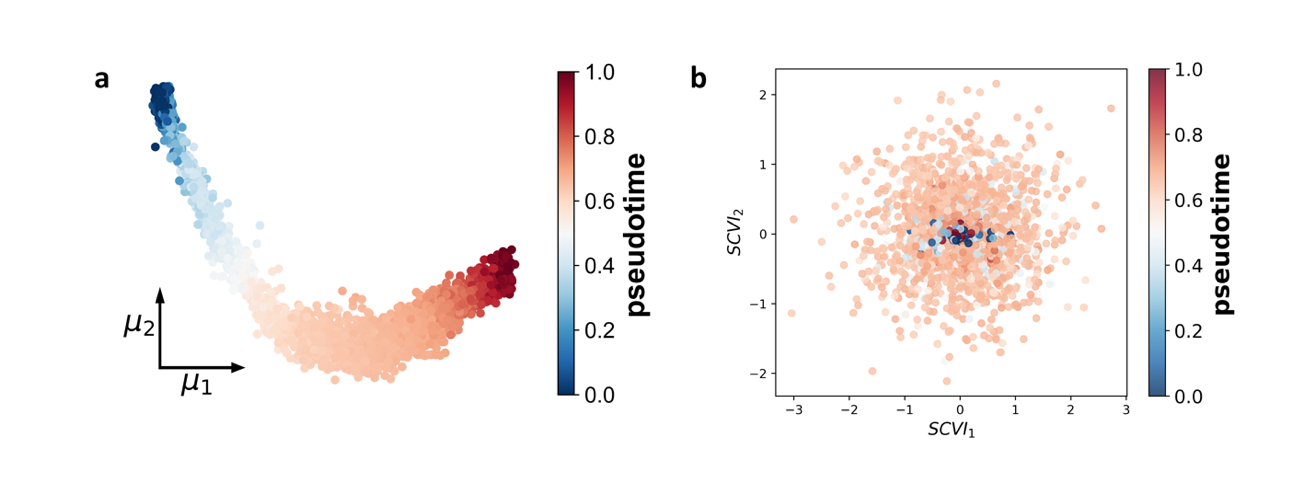

**Figure S5** Comparison of embeddings of SCIM (a) and scVI (b) on dentate gyrus neurogenesis data.

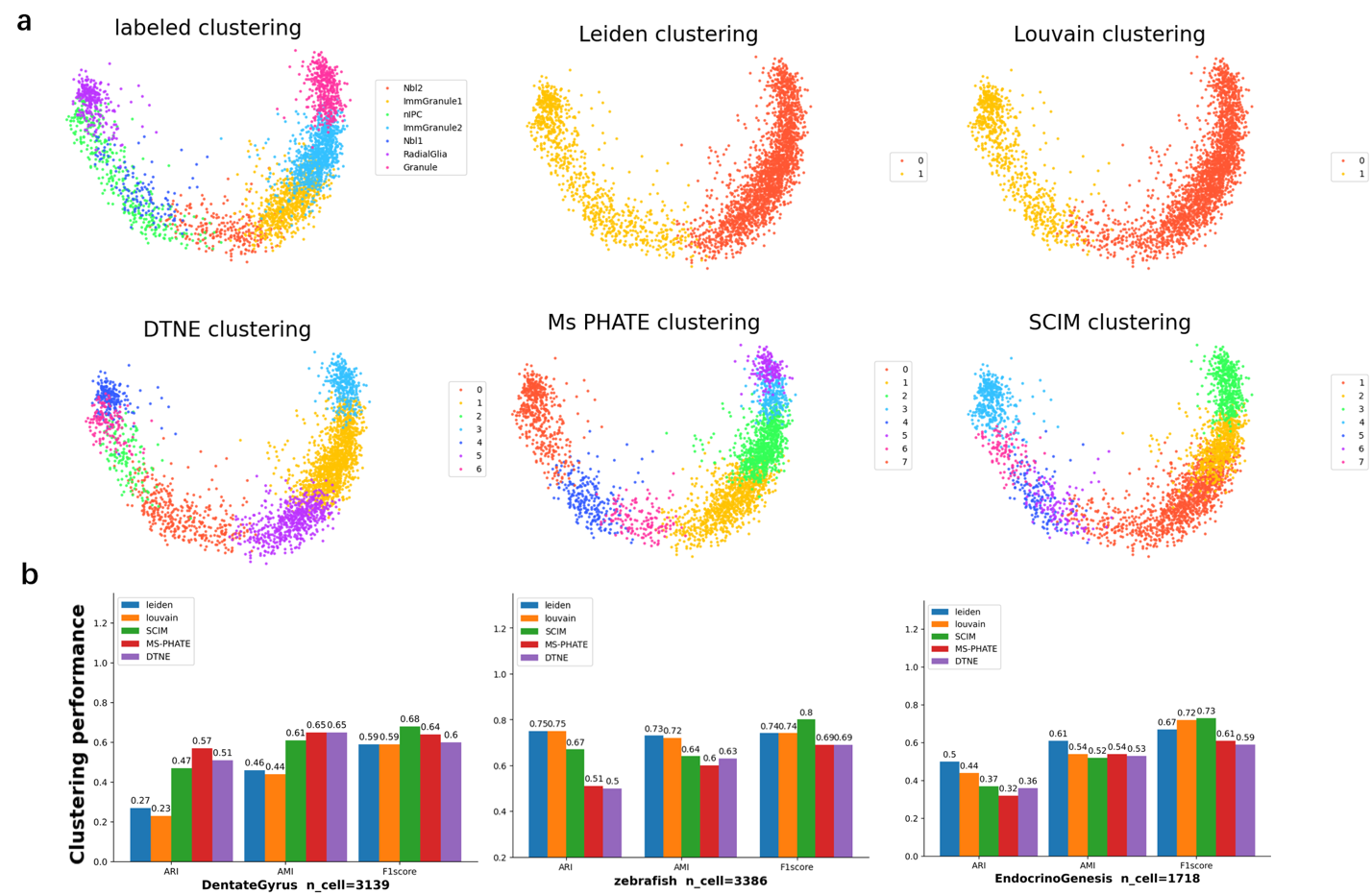

**Figure S6** Comparison between SCIM clustering method and other clustering methods.

(a) Clustering on dentate gyrus data with different methods

(b) Evaluation of five different clustering methods on three single-cell datasets with ARI, AMI and F1-score as performance metrices.

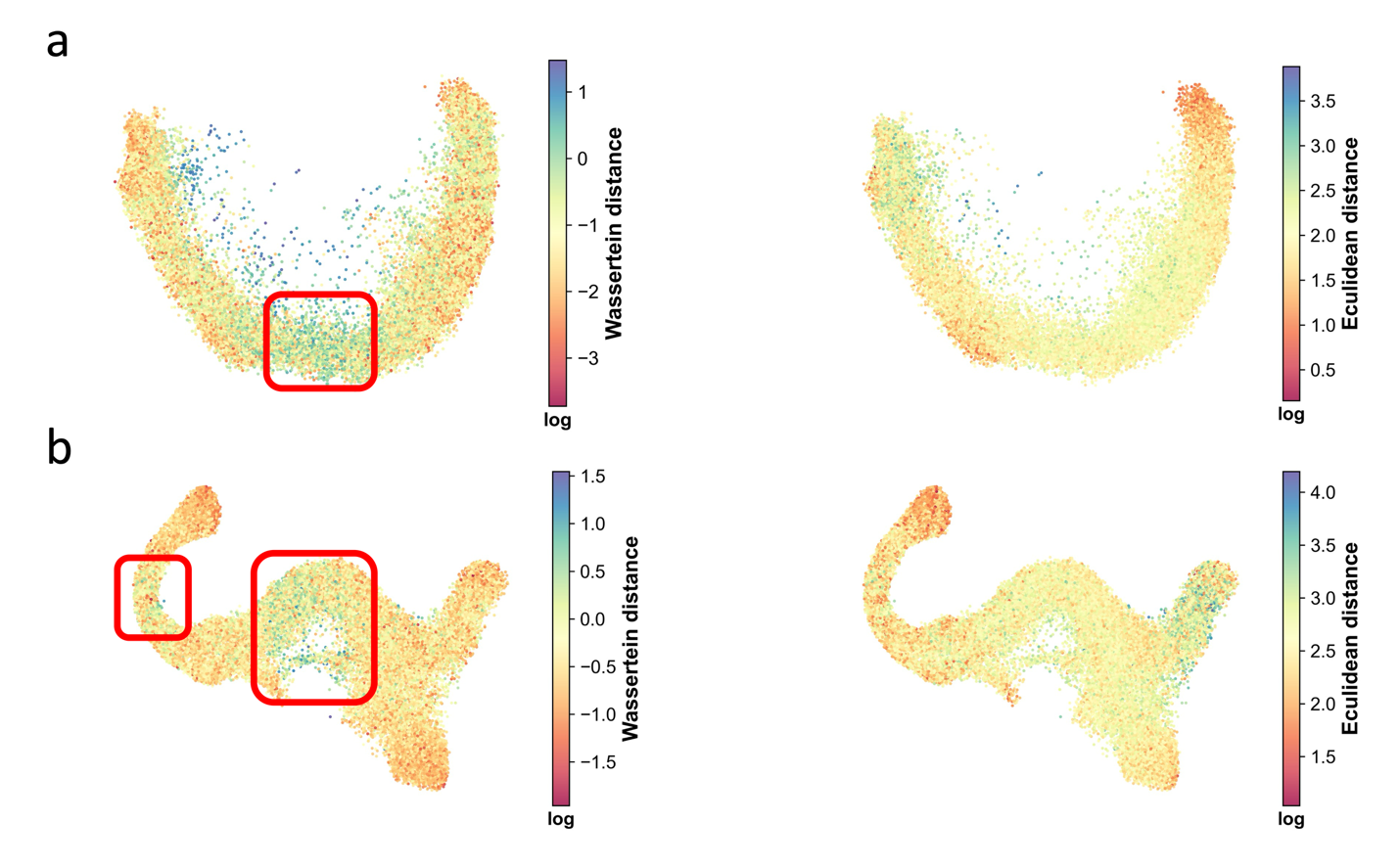

**Figure S7** Low CRC in transition cells arise from variation of Wasserstein distance instead of Euclidean distance between neighboring cells. For better visualization, log scale of distance is shown. Each dot represents an edge on the cell graph.

(a) Wasserstein distance and Euclidean distance in dentate gyrus neurogenesis.
(b) Wasserstein distance and Euclidean distance in pancreatic endocrinogenesis.

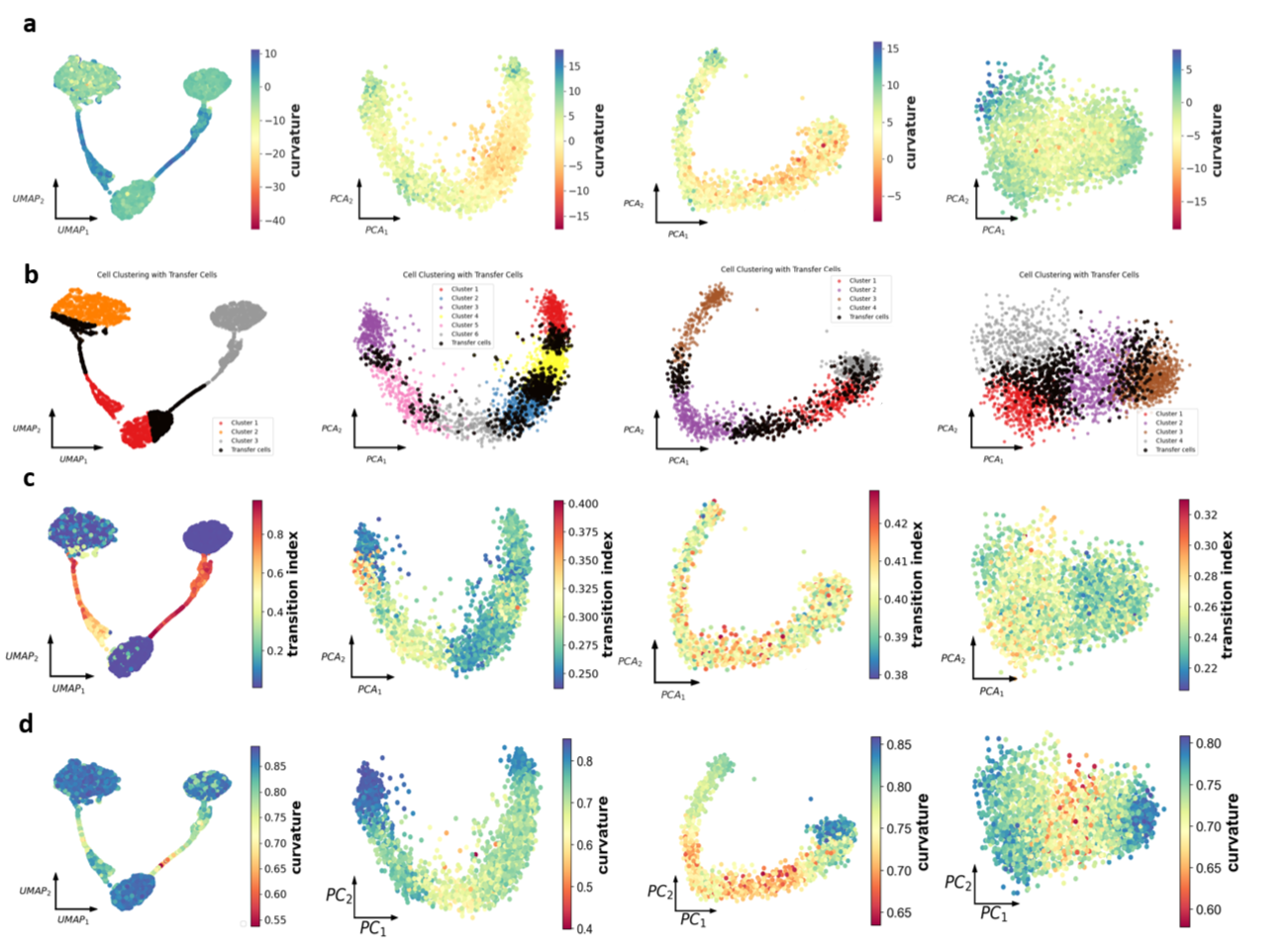
 **Figure S8** Comparison of different transition cells identification methods

(a) CRC calculated via scGeom.
(b) Identification of transition cell with scTite. Transition cells are colored in black.
(c) Transition index calculated via CellTran.
(d) CRC calculated via SCIM.

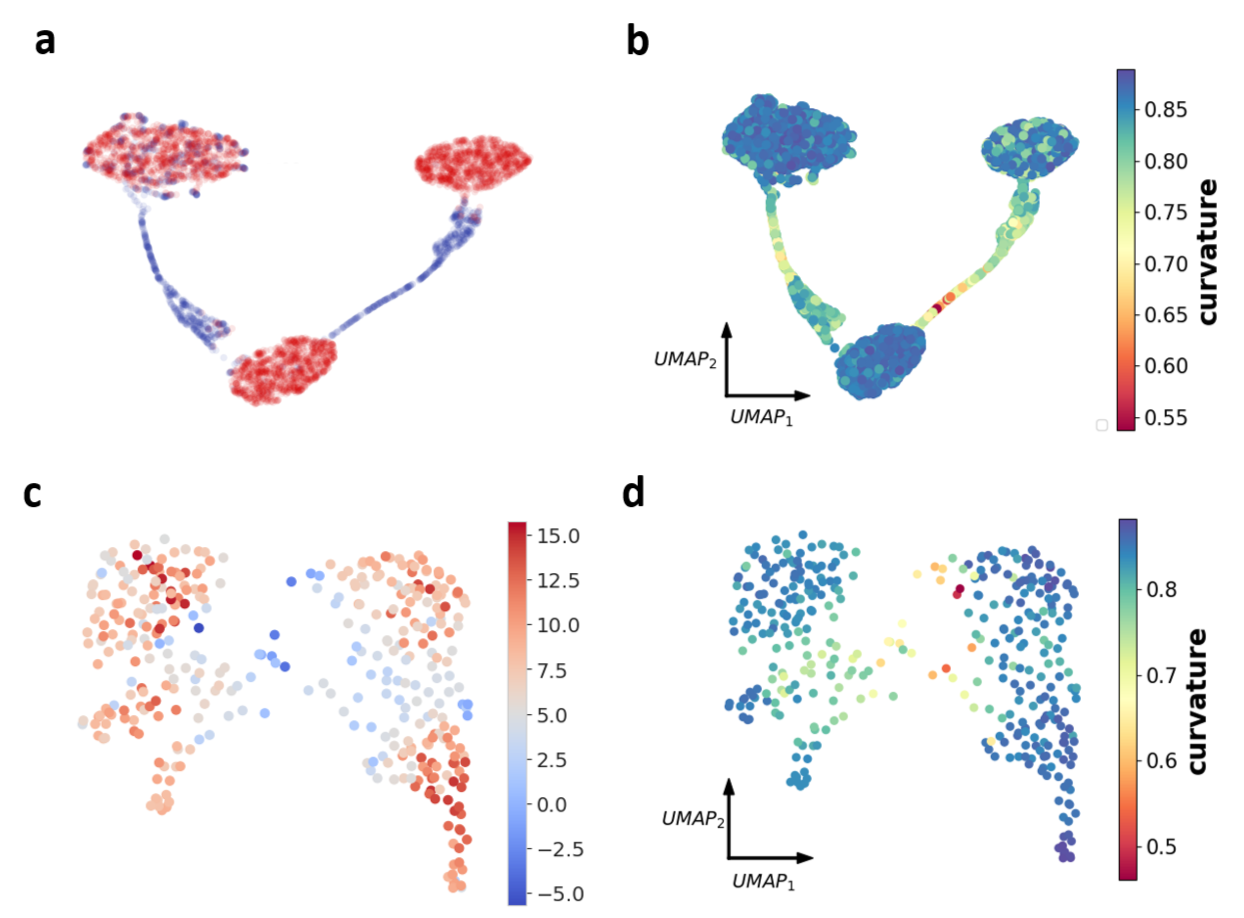

**Figure S9** SCIM on CellTran simulated data and Olsson dataset in scGeom.

(a) Transition cells and stable cells annotation on CellTran simulated data (stable cells are colored in red, while transition cells are in blue)

(b) CRC of CellTran simulated data calculated with SCIM.

(c) CRC of Olsson data calculated with scGeom. Transition cells are Multi-lineage and Mono-interm.

(d) CRC of Olsson data calculated with SCIM.

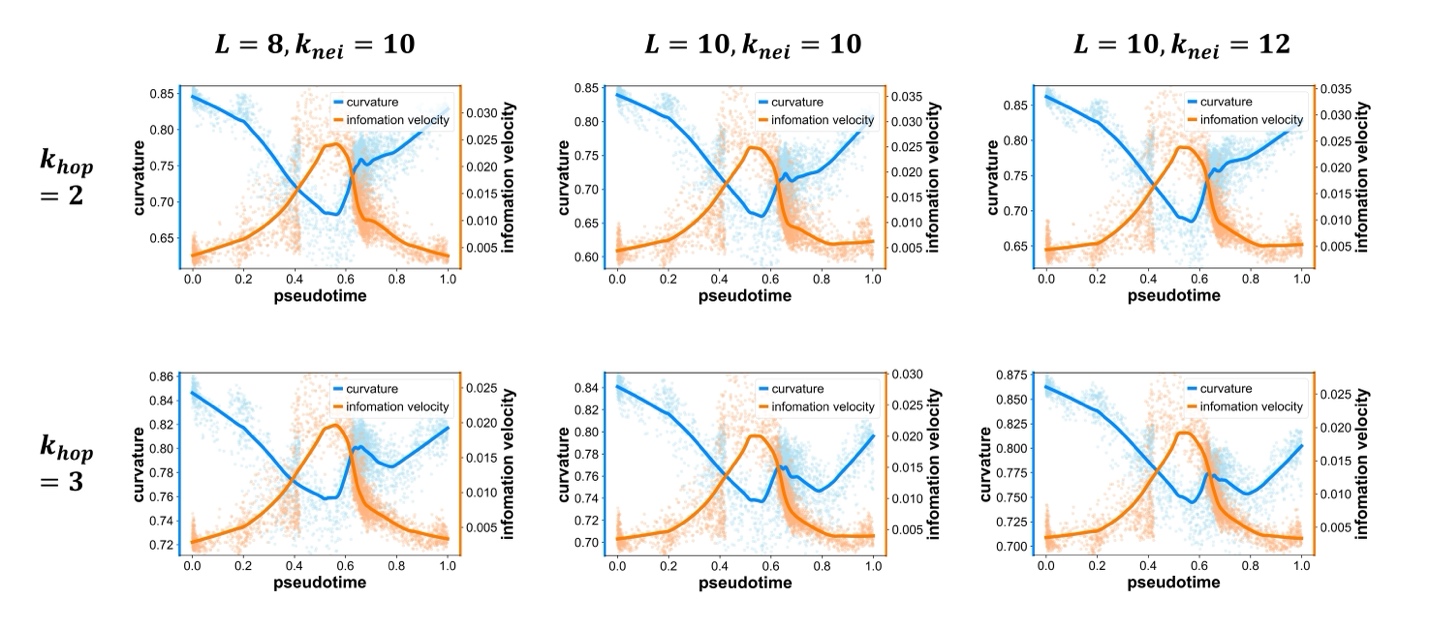

**Figure S10** CRC and information velocity variation in dentate gyrus neurogenesis (with respect to pseudo-time) that are computed with varied values of parameters including latent dimensions (*L*), $k_{nei}$ and $k_{hop}$.

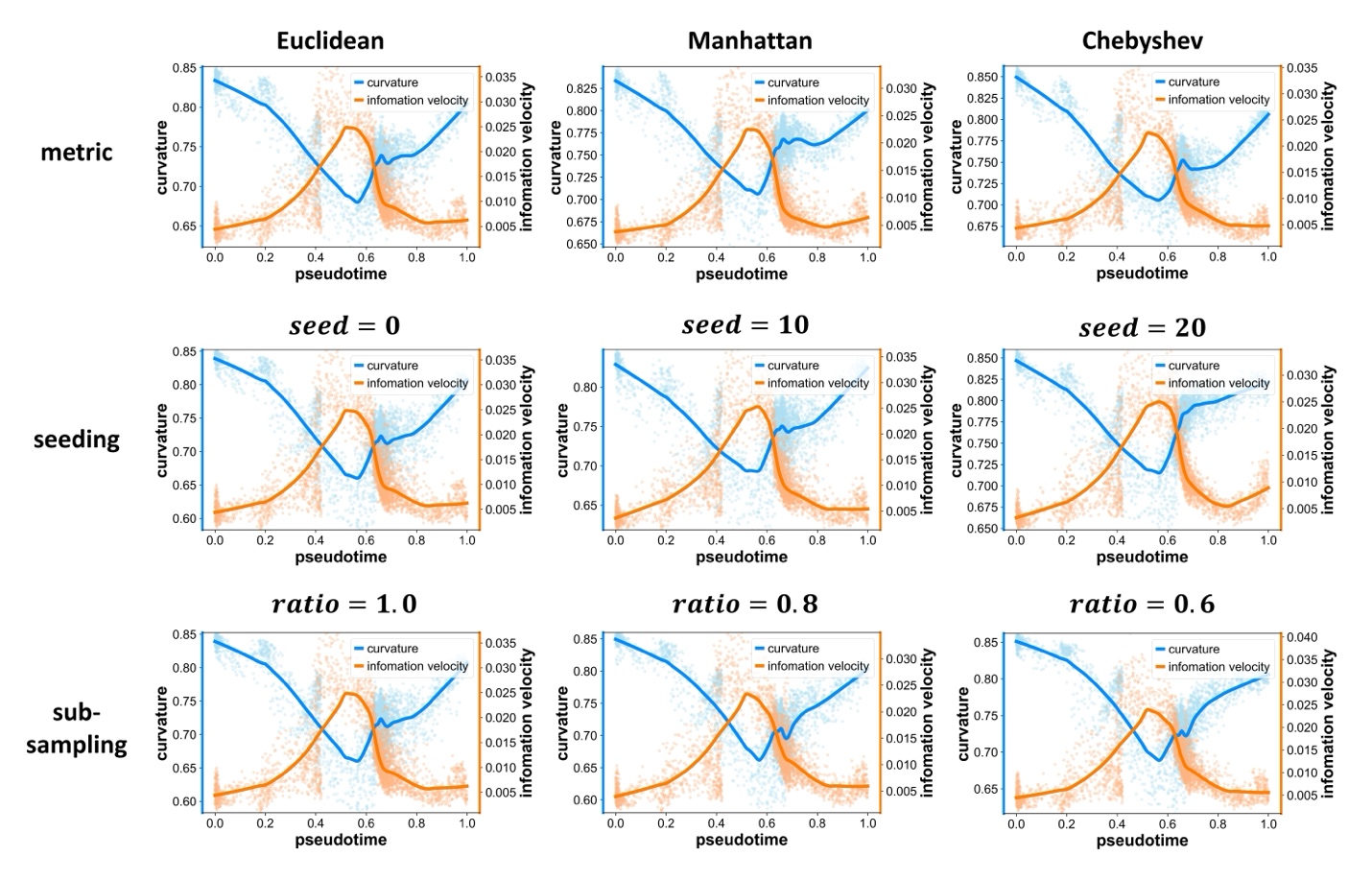

**Figure S11** CRC and information velocity variation in dentate gyrus neurogenesis (with respect to pseudo-time) that are computed with varied values of parameters including different metric of constructing kNN cell graph (top), different seeding values for initialization of training (middle) and different ratios of sub-sampling from the original dataset (bottom).

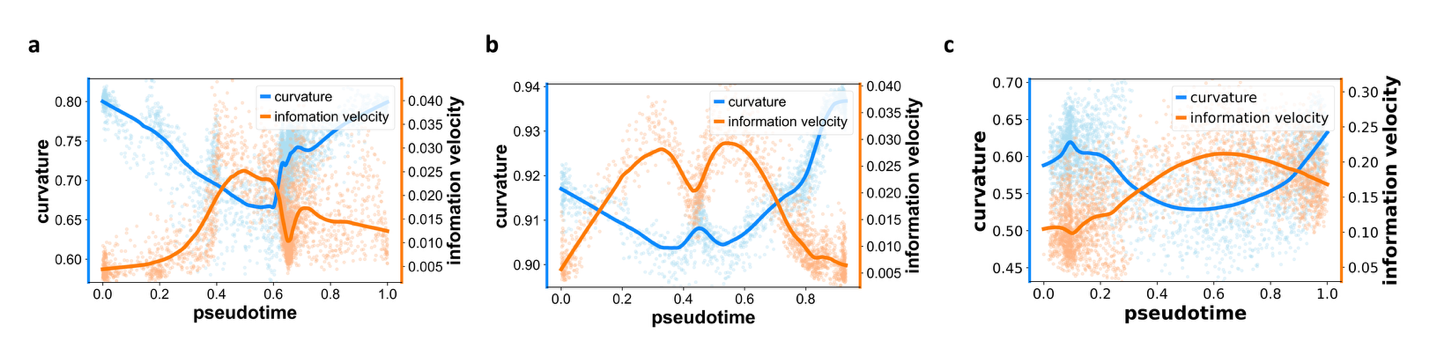

**Figure S12** Analyses of datasets processed with *DUBstepR* method. Correspondence of low-CRC value and high information velocity in dentate gyrus neurogenesis (a), pancreatic endocrinogenesis (b) and EMT of A549 cells (c) (with respect to pseudo-time). The lines are fitted with LOWESS. The pseudotime of pancreatic endocrinogenesis is the palantir pseudotime adapted from the original data file.

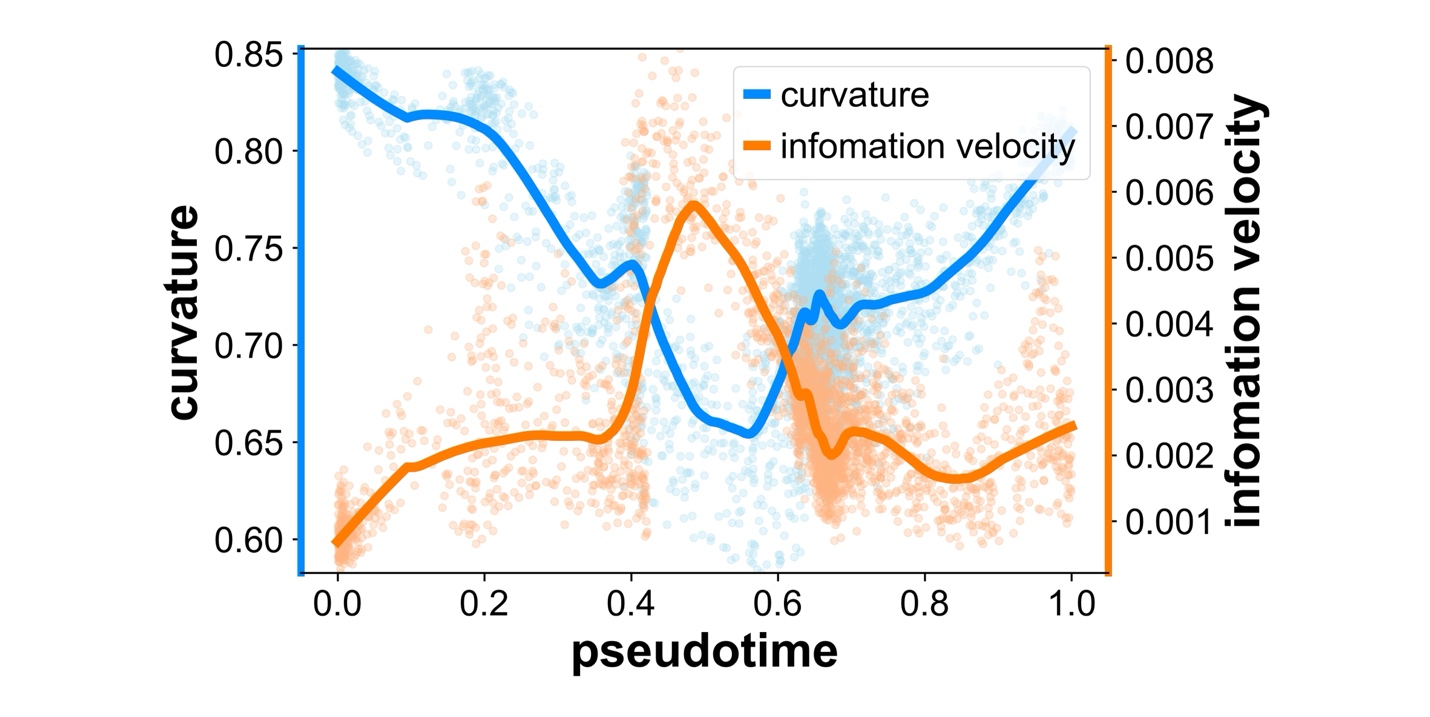

**Figure S13** Correspondence between CRC and information velocity calculated within the NMF coordinates.

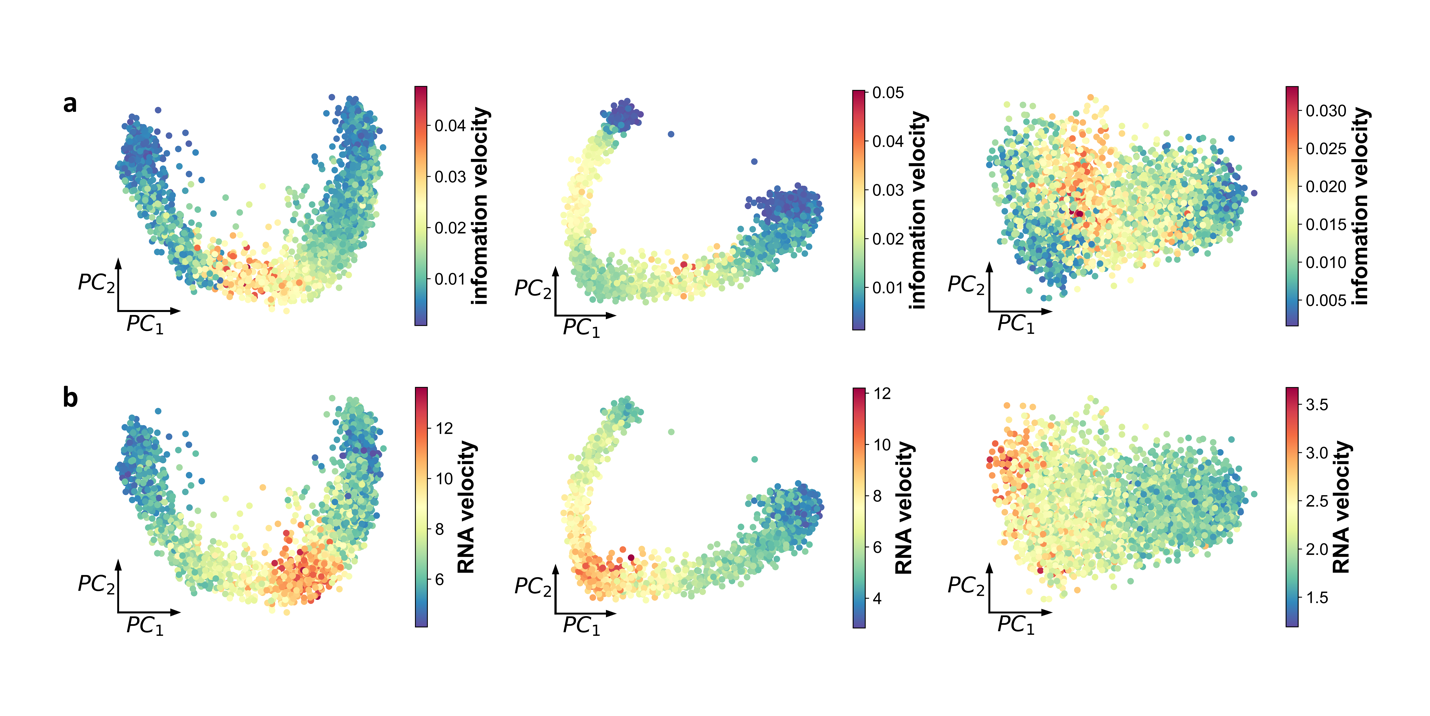

**Figure S14** Comparison between the distribution of information velocity (a) and norm of RNA velocity (b) in dentate gyrus neurogenesis (left), pancreatic endocrinogenesis (middle), and EMT of A549 cells induced by TGF-β (right).

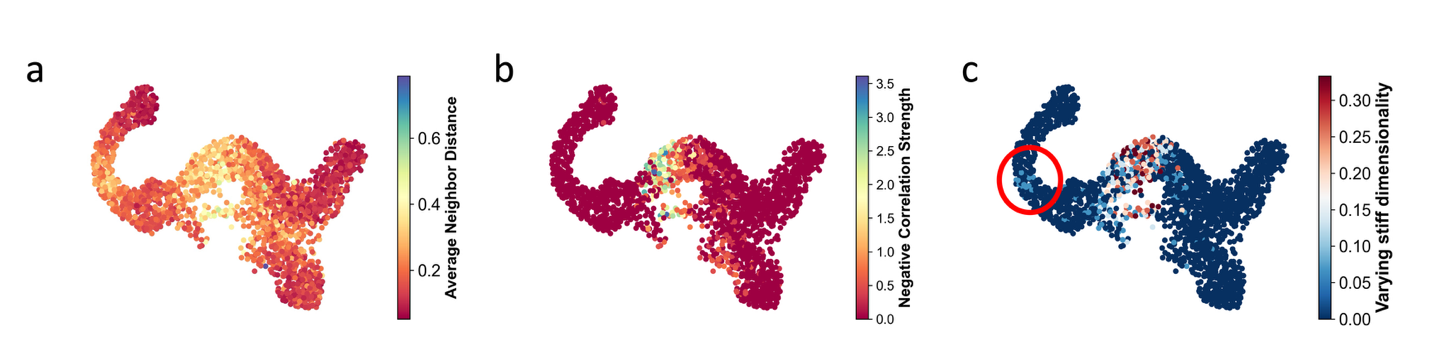

**Figure S15** Variation of stiff directions around branch points.

(a) Counterpart of latent embedding dimension *i* in averaged Wasserstein distance between neighbor cells.
(b) Negative correlation strength of Fisher information of latent embedding dimension *i* in neighbors of each cell.
(c) The ratio of number of varying stiff dimensionality to *L* peaks around the branch points (*L*=15).

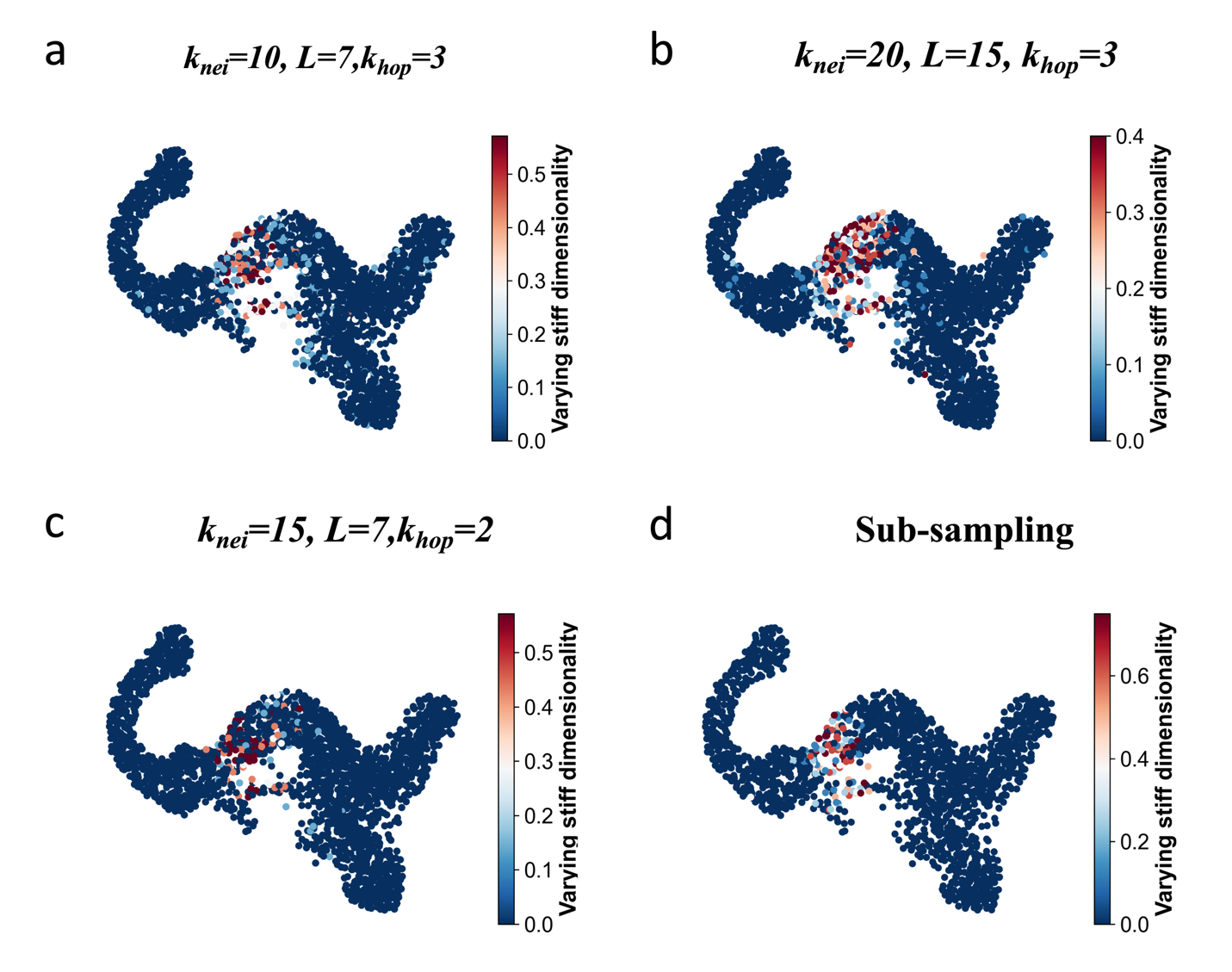

Figure S16 Robustness of varying stiff dimensionality under different parameter settings (a-c) and random sub-sampling (d).

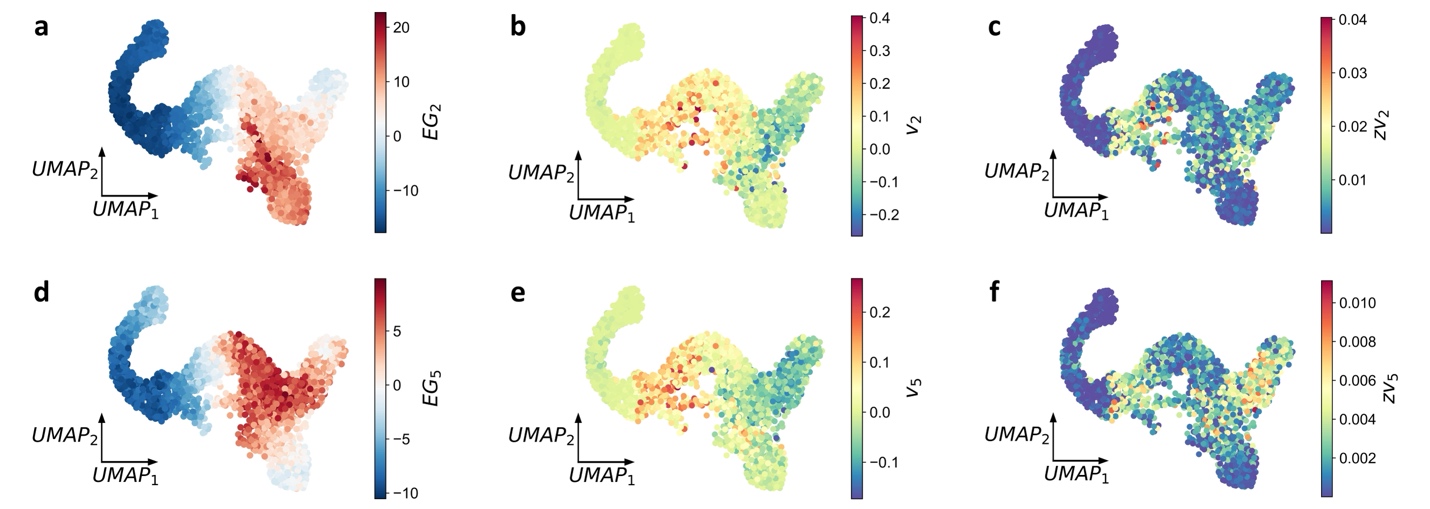

**Figure S17** Time derivative and information velocity of eigengene in the branching process of pancreatic endocriongenesis (including α-cell branch and b-cell branch).

(a) The distribution of eigengene (EG2) that corresponding with α-cell branch (bottom branch) across the whole cell population.

(b) The distribution of time derivative (parameter velocity) of eigengene (EG2) shows a peak around the branch point.

(c) The distribution of information velocity of eigengene (EG2) shows sharp increase at the branch point.

(d) Same as that in panel (a) except for b-cell branch (top branch).

(e) Same as that in panel (b) except for b-cell branch.

(f) Same as that in panel (c) except for b-cell branch.
